# MHC class I and MHC class II reporter mice enable analysis of immune oligodendroglia in mouse models of multiple sclerosis

**DOI:** 10.1101/2022.09.27.509669

**Authors:** Em P Harrington, Riley B Catenacci, Matthew D Smith, Dongeun Heo, Cecilia E Miller, Keya R Meyers, Jenna Glatzer, Dwight E Bergles, Peter A Calabresi

**Author notes:** Corresponding author electronic address. co-first author (Em P Harrington, Riley B Catenacci).

## Abstract

Oligodendrocytes and their progenitors upregulate MHC pathways in response to inflammation, but the frequency of this phenotypic change is unknown and the features of these immune oligodendroglia are poorly defined. We generated MHC class I and II transgenic reporter mice to define their dynamics in response to inflammatory demyelination, providing a means to monitor MHC activation in diverse cell types in living mice and define their roles in aging, injury and disease.

## Introduction

Single-cell and single-nucleus RNA sequencing has revealed that some oligodendroglia in both mouse inflammatory models^1,2^ and human multiple sclerosis (MS)^3–5^ express transcripts associated with major histocompatibility complex (MHC) antigen presenting and processing pathways. These immune oligodendrocyte precursor cells (iOPCs) and oligodendrocytes (iOLs) have been detected in Alzheimer’s disease^6^ and viral infection models^7–9^, and can be induced by exposure to interferon-*γ* (IFN-*γ*), suggesting that some oligodendroglia undergo this distinct phenotypic change in response to inflammation. The role of these immune oligodendroglia is unknown^10^, but their presence raises the possibility that oligodendroglia may present antigens to T cells, be subject to cytotoxic CD8 T cell-mediated death^11^ and perpetuate the immune response through release of cytokines and interactions with CD4 T cells. Oligodendroglial death or inflammation could contribute to impaired remyelination seen in MS and other progressive diseases^12-14^. Further exploration of the spatial and temporal dynamics of iOPCs/iOLs have been limited by their relative rarity and our inability to identify which cells have transformed in living tissue. To enable detection of which cells upregulate MHC pathways *in vivo*, we generated two novel MHC I and MHC II reporter mouse lines that express tdTomato when these pathways are activated.

## Results

### MHC reporter mice are a reliable readout of MHC protein expression

MHC class II chaperone invariant chain (CD74 or Ii) and MHC class I component beta-2-microglobulin (B2m) are required components of the antigen processing/presentation machinery that are expressed by a subset of oligodendroglia in mouse inflammatory models^1^ and human MS brain^3,4^. To facilitate identification of these cells, we used CRISPR/Cas9 mediated gene editing to replace the stop codon of these genes with a P2A-TdTomato-WPRE-pA sequence (Figure 1A-C and Figure 1- figure supplement 1), generating *B2m-TdTomato* (*B2m-TdT*) and *CD74-TdTomato* (*CD74-TdT*) reporter mice without disrupting expression of the endogenous genes (Figure 1- figure supplement 2).

**Figure 1.**
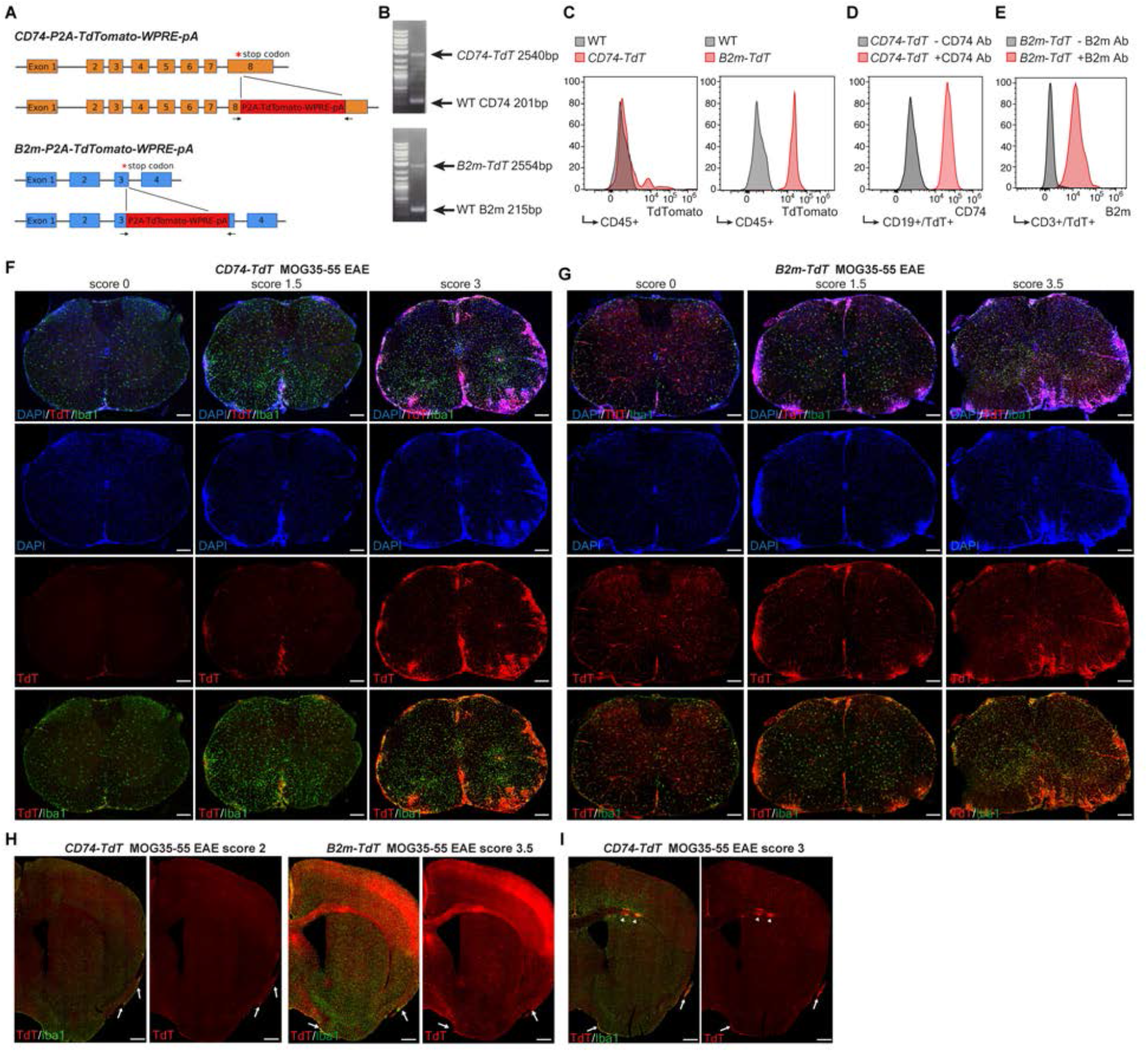
MHC reporter mice validation and MOG_35-55_ peptide immunized EAE brain and spinal cord TdTomato reporter expression. (**A**). Stop codons of CD74 and B2m are replaced by P2A-TdTomato-WPRE-pA construct to generate *CD74-TdT* and *B2m-TdT* reporter mice, respectively. Figure made in Biorender.com. (**B**). Genotyping of reporter mice with primers (indicated by arrows in panel A) spanning reporter construct insertion site. (**C**). Flow cytometry of endogenous TdT expression in peripheral blood CD45+ cells for phenotyping reporter and wild-type mice. (**D**). Flow cytometry of TdT+CD19+ B cells stained with and without CD74 antibody from *CD74-TdT* spleen. (**E**). Flow cytometry of TdT+CD3+ T cells stained with and without B2m antibody from *B2m-TdT* spleen. (**F**). Representative images of *CD74-TdT* reporter MOG_35-55_ EAE spinal cord. Endogenous TdT expression is restricted to the meninges in pre-clinical score 0 mice and with increasing clinical score is prominent within Iba1+ reactive clusters and DAPI hypercellular lesion areas. Scale bars, 200μm. (**G**). Representative images of *B2m-TdT* reporter MOG_35-55_ EAE spinal cord. Endogenous TdT expression is present in the vasculature and Iba1+ microglia in pre-clinical score 0 animals and TdT demonstrates diffuse expression within lesion areas and more notable parenchymal expression with increasing clinical score. Scale bars, 200μm. (**H**). Representative images of *CD74-TdT* and *B2m-TdT* reporter MOG_35-55_ EAE brain. TdT endogenous expression is found in meningeal clusters (arrows). Scale bars, 500μm. (**I**). Representative images of *CD74-TdT* EAE brain with TdT clusters the corpus callosum (arrowheads) and brain parenchyma. Scale bars, 500μm.

**Figure 1- figure supplement 1.**
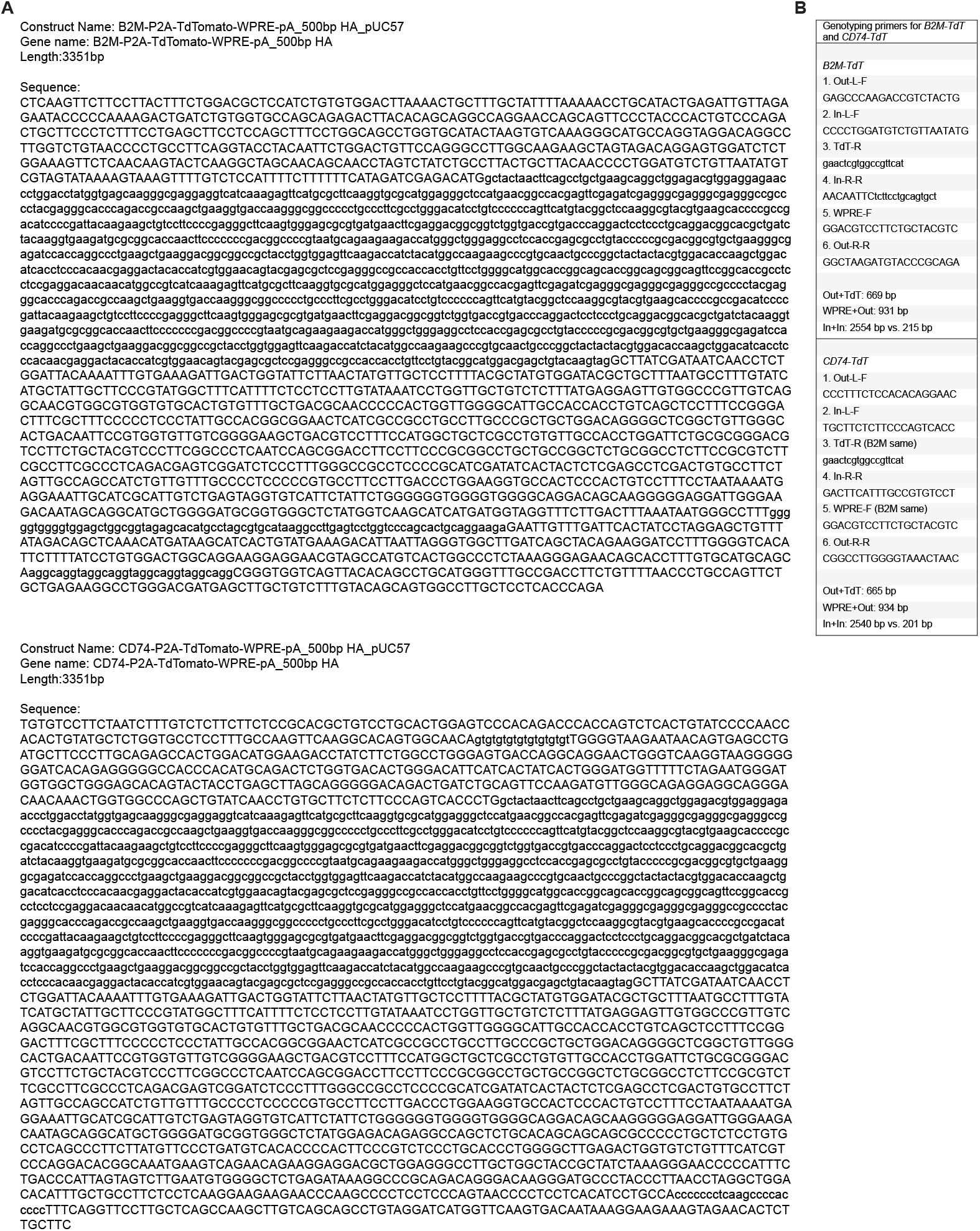
Generation of *B2m-TdT* and *CD74-TdT* reporter animals. (**A**) CD74-P2A-TdTomato-WPRE-pA and B2M-P2A-TdTomato-WPRE-pA construct sequences with 500bp homology arms. First uppercase sequence 500bp homology arm followed by lowercase sequence P2A-TdTomato and upper-case sequence WPRE-pA and 500bp homology arm. (**B**). Primers for genotyping *B2m-TdT* and *CD74-TdT* reporter animals.

**Figure 1- figure supplement 2.**
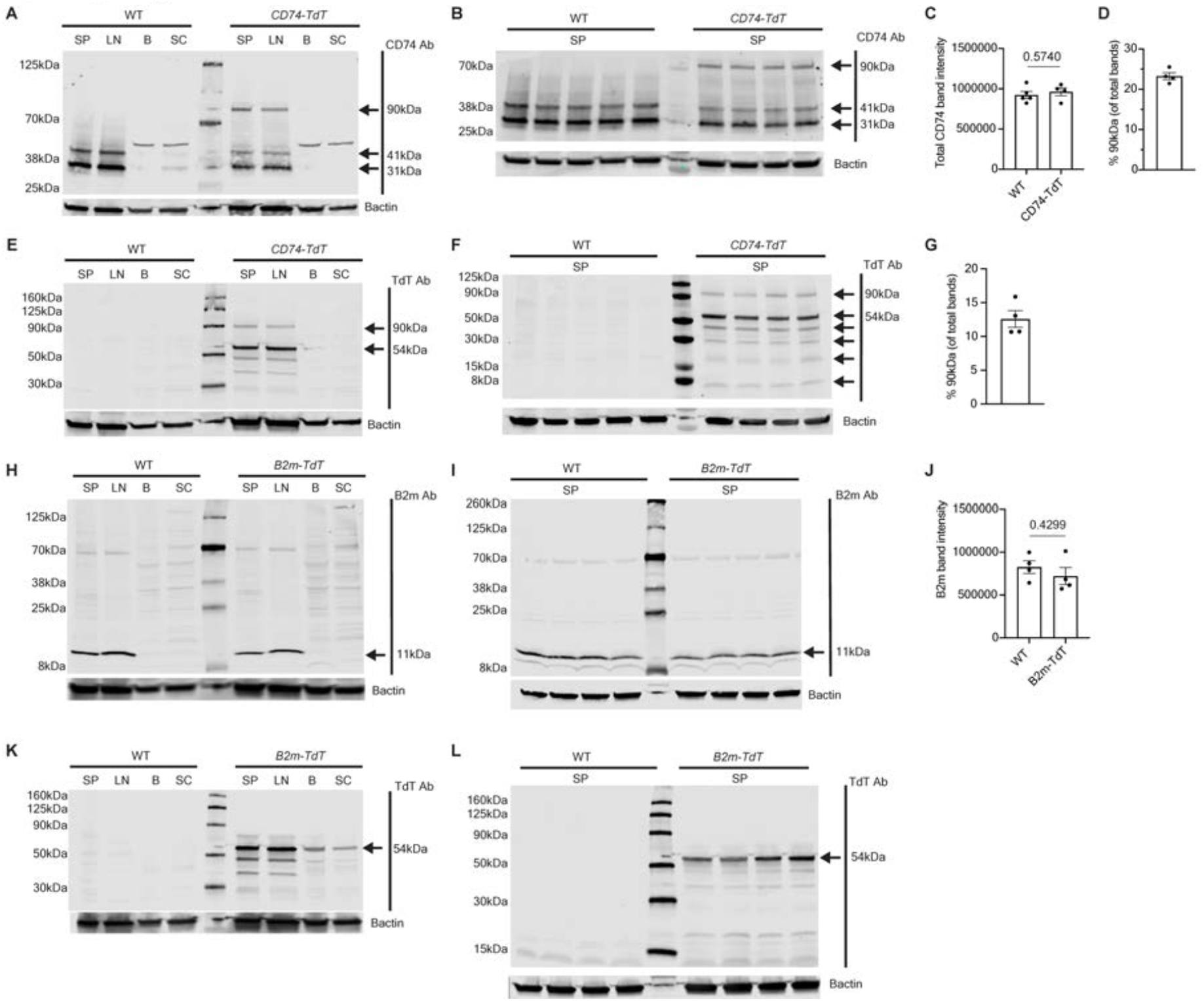
Western blots of *B2m-TdT* and *CD74-TdT* reporter and WT animals. (**A**). *CD74-TdT* and WT spleen, lymph node, brain, and spinal cord tissue probed with CD74 antibody. Expected p31/p41 CD74 isoforms and 90kDa *CD74-TdT* fusion protein are present in *CD74-TdT* spleen and lymph nodes, brain with single CD74 isoform above p41 isoform. (**B**). *CD74-TdT* fusion protein (90kDa) present in *CD74-TdT* spleen probed for CD74 antibody. (**C**). Quantification of total CD74 protein (combined 31, 41, 90 kDa band) intensity in WT and *CD74-TdT* spleen, n = 4 mice/group. Unpaired t-test. (**D**). Quantification of percentage of 90kDa *CD74-TdT* fusion protein in *CD74-TdT* spleen, n = 4 mice. (**E**). *CD74-TdT* and WT spleen, lymph node, brain, and spinal cord tissue probed with TdT antibody. TdT protein (54kDa) and CD74-TdT fusion protein (90kDa) present in *CD74-TdT* spleen and lymph node. (**F**). *CD74-TdT* fusion protein (90kDa) present in *CD74-TdT* spleen probed for TdT antibody. (**G**). Quantification of percentage of 90kDa *CD74-TdT* fusion protein in *CD74-TdT* spleen, n = 4 mice. (**H**). *B2m-TdT* and WT spleen, lymph node, brain, and spinal cord tissue probed with B2m antibody with expected 11kDa B2m isoform protein band in spleen and lymph node. (**I**). WT and *B2m-TdT* spleen probed for B2m antibody. (**J**). Quantification of total B2m protein band intensity in WT and *B2m-TdT* spleen, n = 4 mice/group. Unpaired t-test. (**K**). *B2m-TdT* and WT spleen, lymph node, brain, and spinal cord tissue probed with TdT antibody with expected 54kDa TdT band in reporter tissue. (**L**). *B2m-TdT* and WT spleen probed for B2m antibody with single 54kDa TdT band in *B2m-TdT* spleen. Data represented are means ± s.e.m.; 20μg protein loaded per lane, each lane is an individual animal. B-brain; LN-lymph node; SP-spleen; SC-spinal cord; WT-wild-type. Figure 1- figure supplement 2- source data 1 Original western blot images. Blots are labeled with antibody probe, samples and upper left letter corresponds to letter depicted in Figure 1- figure supplement 2. Figure 1- figure supplement 2- source data 2 Data from analysis of western blot band intensities. Sheets are labeled with letter corresponding to data in Figure 1- figure supplement 2.

To determine if these transgenes accurately report transcriptional activation of MHC components, we examined splenic immune cells known to express MHC class I and II. In *CD74-TdT* reporter mice, TdT expression was highest in professional antigen presenting cells, such as dendritic and B cells that were immunoreactive to CD74 (Figure 1D and Figure 1- figure supplement 3A,B). Independent of MHC class II, CD74 has diverse roles in cell survival, migration and MIF (macrophage migration inhibitory factor) cytokine signaling^15,16^. Thus, we determined the relationship between TdT and MHC class II expression using I-A/I-E MHC antibodies (Figure 1- figure supplement 3C-E). Cells that exhibited intermediate TdT fluorescence were also negative or weakly immunoreactive for I-A/I-E expression (Figure 1- figure supplement 3D), indicating that the level of TdT expression correlates well with MHC class II receptor expression. In *B2m-TdT* reporter mice, TdT was ubiquitously expressed in all immune cells analyzed in the spleen and TdT fluorescent cells were immunoreactive to B2m (Figure 1E).

**Figure 1- figure supplement 3.**
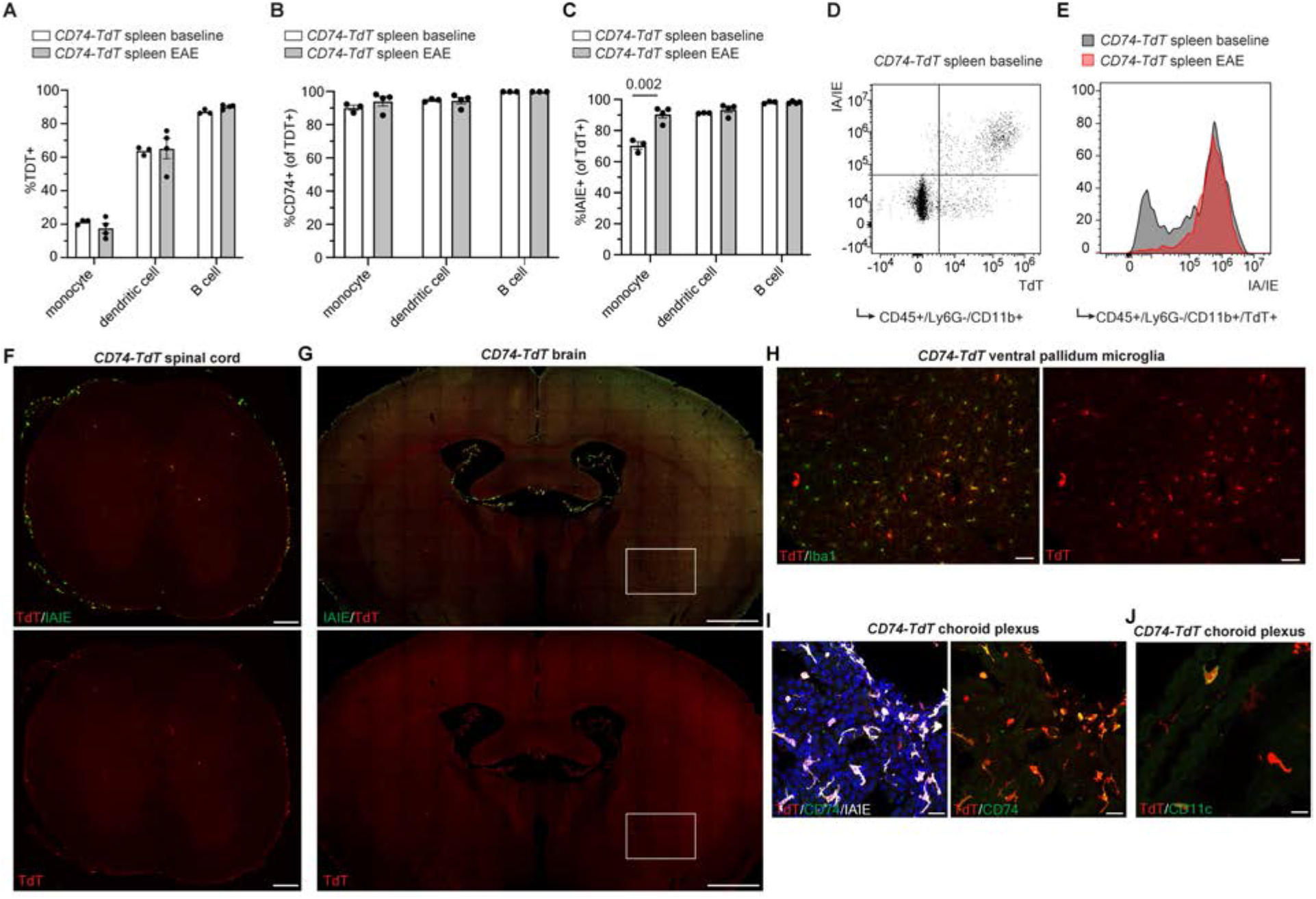
CD74-TdT reporter baseline and EAE expression. (**A**). Flow cytometry of spleen endogenous *CD74-TdT*+ populations at baseline and MOG_35-55_ EAE, n = 3-4 mice/group. (**B**). Flow cytometry of spleen endogenous TdT co-expression with CD74 antibody at baseline and MOG_35-55_ EAE, n = 3-4 mice/group. (**C**). Flow cytometry of spleen endogenous TdT co-expression with MHC class II IA/IE antibody in non-immunized and MOG_35-55_ EAE, n = 3-4 mice/group. Unpaired t-test. (**D**). Representative flow cytometry plot of TdT and IA/IE expression in monocytes in baseline spleen with IA/IE negative/TdT intermediate population. (**E**). Flow cytometry representative histogram of IA/IE expression in TdT+CD11b+ monocytes, with TdT+ IA/IE-negative population not present in immunized spleen. (**F**). Representative images of *CD74-TdT* endogenous reporter expression in baseline adult spinal cord stained for IA/IE. Scale bars, 200μm. (**G**). Representative images of *CD74-TdT* endogenous reporter expression in baseline adult brain stained for IA/IE. Scale bars, 1mm. (**H**). Representative images of TdT expression in subset of Iba1+ microglia in ventral pallidum (magnification of box in **G**). Scale bars, 50μm. (**I**). Representative confocal images of baseline adult brain choroid plexus endogenous TdT stained for CD74 and IA/IE. Scale bars, 20μm. (**J**). Representative confocal images of baseline choroid plexus endogenous TdT staining of CD11c+ dendritic cells. Scale bars, 20μm. Data represented are means ± s.e.m. Figure 1- figure supplement 3- source data 1. Data from analysis of flow cytometry depicted in Figure 1=- figure supplement 3. Sheets are labeled with letter corresponding to data in Figure 1- figure supplement 3.

In the adult central nervous system (CNS), TdT expression in *CD74-TdT* mice was most prevalent in the meninges and choroid plexus, but was also observed within some microglia in the ventral brain (Figure 1- figure supplement 3F-J). In *B2m-TdT* mice, TdT expression was prominent in endothelial cells and microglia throughout the brain (Figure 1- figure supplement 4A-E). In addition, some neurons in the cerebellum, septum, hippocampus and cortex expressed TdT (Figure 1- figure supplement 4F-J). B2m transcripts are upregulated in Cux2 expressing cortical layer II/II pyramidal neurons in human MS, which are susceptible to neurodegeneration^3^. Expression of MHC class I by these neurons may allow them to present antigen and render them more prone to death in inflammatory conditions.

**Figure 1- figure supplement 4.**
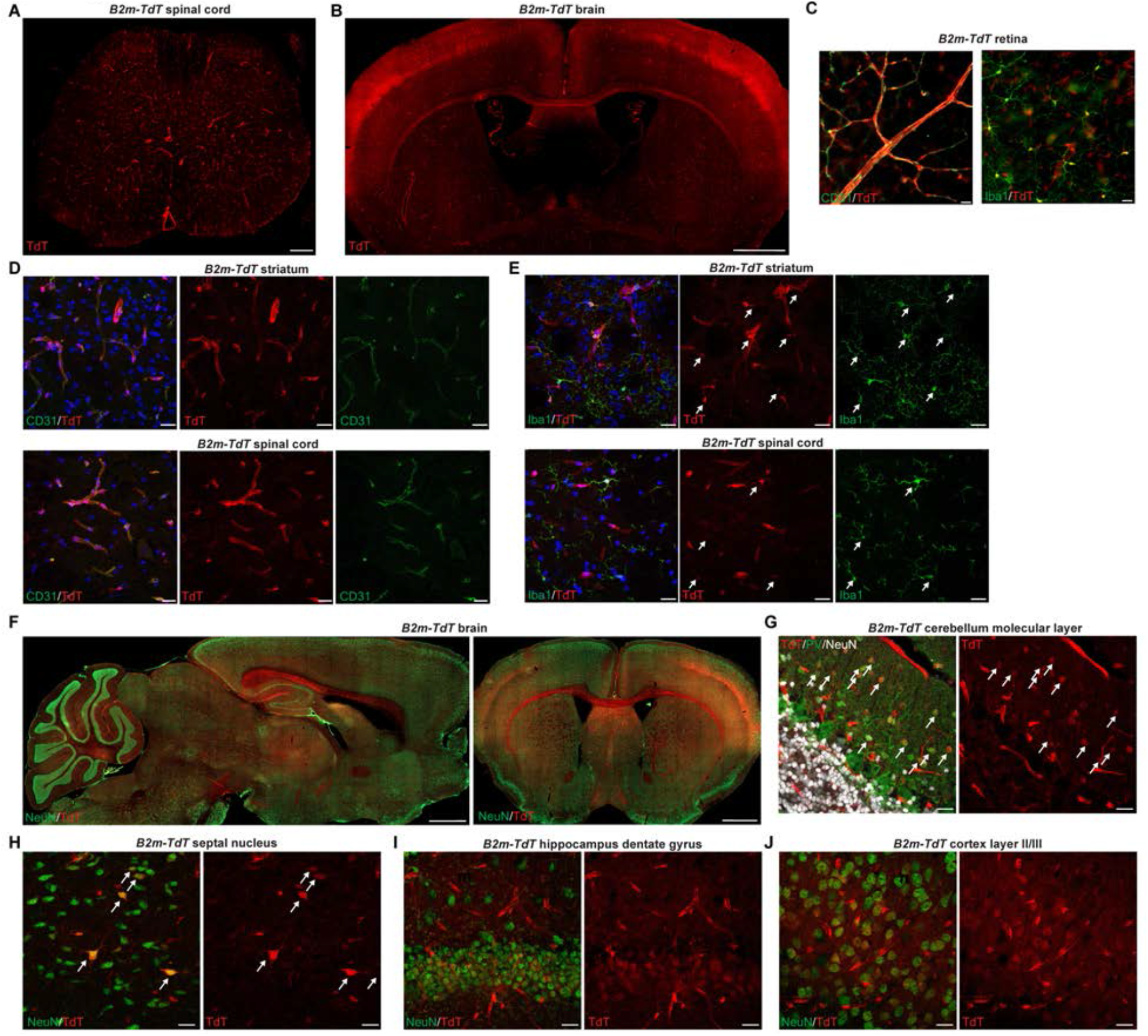
B2m-TdT reporter baseline expression. (**A**). Representative images of *B2m-TdT* endogenous reporter expression in baseline adult spinal cord. Scale bars, 200μm. (**B**). Representative images of *B2m-TdT* endogenous reporter expression in baseline adult brain.Scale bars, 1mm. (**C**). Representative images of flat mount adult *B2m-TdT* retina with TdT expression on CD31+ endothelial cells and Iba1+ microglia in retina. Scale bars, 20μm. (**D**). Representative confocal imaging of striatum and spinal cord endogenous TdT expression on CD31+ endothelial cells. Scale bars, 20μm. (**E**). Representative confocal imaging of striatum and spinal cord endogenous TdT expression on Iba1+ microglia with TdT+Iba1+ cell bodies indicated with arrows. Scale bars, 20μm. (**F**). Representative images of *B2m-TdT* endogenous reporter expression in sagittal brain and coronal brain stained with NeuN. Scale bars, 1mm. (**G**). Representative confocal imaging of endogenous TdT+ parvalbumin PV+ neurons, indicated by arrows, in cerebellar molecular layer. Scale bars, 20μm. (**H**). Representative confocal imaging of septal nucleus with scattered endogenous TdT+ neurons. Scale bars, 20μm. (**I**). Representative confocal imaging of hippocampal dentate gyrus endogenous TdT+ granule cells. Scale bars, 20μm. (**J**). Confocal imaging of endogenous TdT+ neurons in cortex layer II/III. Scale bars, 20μm. Data represented are means ± s.e.m.

### MHC reporter expression is induced in EAE

To evaluate CNS activation of MHC pathways in distinct cell types under inflammatory conditions, we induced experimental autoimmune encephalitis (EAE) by MOG_35-55_ peptide immunization. Reporter mice subjected to MOG_35-55_ EAE were sacrificed with EAE presentation ranging from pre-clinical (score 0), tail and hindlimb weakness (score 1-2.5) and complete hindlimb paralysis (score 3-4). With increasing EAE clinical score, there was a concomitant increase in TdT in spinal cord lesion areas in both *CD74-TdT* (Figure 1F) and *B2m-TdT* animals (Figure 1G). TdT was also present in EAE ventral brain meningeal cell clusters in *CD74-TdT* and *B2m-TdT* animals (Figure 1H), and corpus callosum cell clusters in *CD74-TdT* mice (Figure 1I). Infiltrating myeloid cells and microglia exhibited TdT expression in brain and spinal cord of *CD74-TdT* mice with EAE (Figure 1- figure supplement 5A,B) and both the proportion of costimulatory molecule-expressing myeloid cells and microglia that expressed TdT, and the level of TdT expression by these cells, were higher compared to non-co-stimulatory molecule expressing cells (Figure 1- figure supplement 5C-F). In *B2m-TdT* mouse brain and spinal cord, the percentage of TdT expressing microglia was not significantly different between M1 or M2 microglia, defined by CD86 and CD206 expression, respectively^17^ (Figure 1- figure supplement 5G-I); however, M1 microglia had significantly higher TdT fluorescence compared to M2 microglia (Figure 1- figure supplement 5J). Together, this analysis highlights the ability of these reporter mice to reliably identify endogenous and infiltrating cells that exhibit MHC upregulation.

**Figure 1- figure supplement 5.**
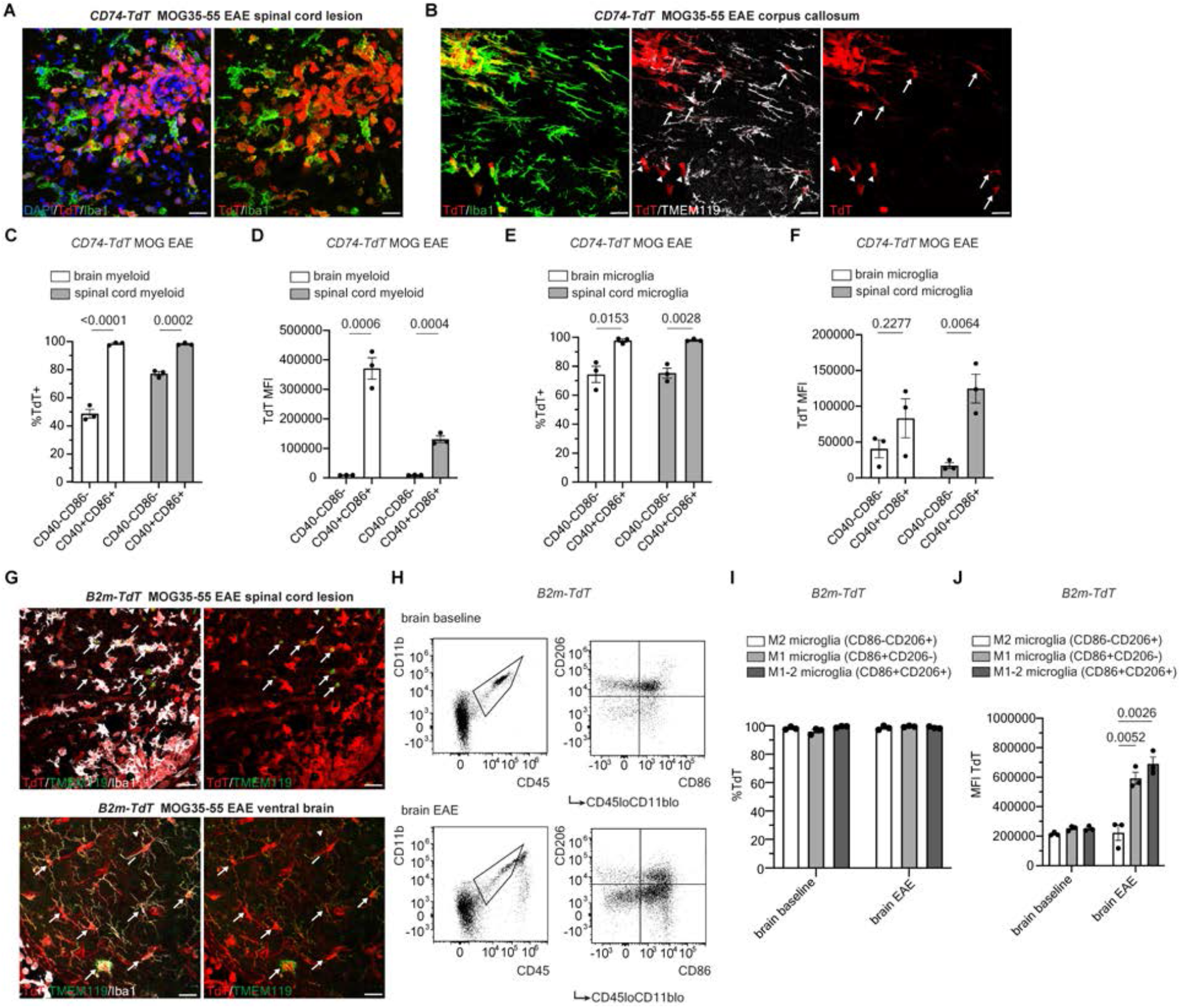
*CD74-TdT* and *B2m-TdT* reporter expression in MOG_35-55_ EAE. **(A)**. Representative confocal imaging of *CD74-TdT* MOG_35-55_ peptide EAE spinal cord lesion with TdT+Iba1+ cells. Scale bars,20µm. **(B)**. Representative confocal imaging of *CD74-TdT* MOG_35-55_ peptide EAE brain corpus callosum lesion with endogenous TdT+ microglia (TdT+Iba1+TMEM119+, arrows) and TdT+ infiltrating myeloid cells (TdT+Iba1+TMEM119−, arrowheads). Scale bars,20µm. **(C)**. Flow cytometry percentage of TdT expressing costimulatory molecule positive (CD40+CD86+) compared to negative (CD40–CD86–) infiltrating myeloid cells (CD45+CD11b+Clec12a+) in CD74–TdT MOG_35-55_ peptide EAE, n = 3 mice/group. Unpaired t-test. **(D)**. Flow cytometry endogenous TdT mean fluorescence intensity (MFI) in TdT+ in costimulatory positive and negative infiltrating myeloid cells (CD45+CD11b+Clec12a+) in CD74–TdT MOG_35-55_ peptide EAE, n = 3 mice/group. Unpaired t-test. **(E)**. Flow cytometry percentage of TdT+ costimulatory positive and negative microglia (CD45+CD11b+Clec12a–) in CD74-TdT MOG_35-55_ peptide EAE, n = 3 mice/group. Unpaired t-test. **(F)**. Flow cytometry of TdT MFI in co-stimulatory positive and negative microglia (CD45+CD11b+Clec12a–) in CD74-TdT MOG_35-55_ peptide EAE, n = 3 mice/group. Unpaired t-test. **(G)**. Representative confocal imaging of B2m-TdT MOG_35-55_ peptide EAE spinal cord and brain TdT+TMEM119+ microglia indicated by arrows. Scale bars, 20µm.**(H)**. Representative gating strategy for M2 (CD86-CD206+), M1 (CD86+CD206-) and M1-2 microglia (CD86+CD206+) in *B2m-TdT* baseline and EAE brain. **(I)**. Flow cytometry percentage of TdT+ M1, M2 and M1-2 microglia in *B2m-TdT* baseline and MOG_35-55_ peptide EAE brain, n = 3 mice/group. **(J)**. Flow cytometry TdT MFI of M1, M2 and M1-2 microglia in *B2m-TdT* baseline and MOG_35-55_ peptide EAE brain, n = 3, mice/group. Unpaired t-test. Data represented are means ± s.e.m. Figure 1- figure supplement 5- source data 1 Data from analysis of flow cytometry depicted in Figure 1- figure supplement 5. Sheets are labeled with letter corresponding to data in Figure 1- figure supplement 5.

### MHC reporter positive immune oligodendroglia correlate with degree of inflammation

We have previously shown that IFN-*γ* is sufficient to induce MHC class I and II expression in oligodendroglia *in vitro*^11^. Consistent with this observation, IFN-*γ* increased TdT expression in Olig2 immunoreactive (+) cells OPC enriched cultures^18^ (Figure 2A,B) and TdT expression was found in Iba1+ microglia and GFAP+ astrocytes from *CD74-TdT* mice treated with IFN-*γ* (Figure 2B). *In vivo*, Olig2+ oligodendroglia were also found to express TdT in both *B2m-TdT* and *CD74-TdT* MOG_35-55_ EAE spinal cord, the main CNS region affected by EAE^19^ (Figure 2C), in accordance with the profound increase in IFN-*γ* observed during EAE. The number of TdT+Olig2+ oligodendroglia was significantly higher in clinically symptomatic compared to pre-symptomatic score 0 animals in both *B2m-TdT* (Pre: 0.9 ± 0.2%, n=4; Post: 6.4 ± 1.7%, n=9, p=0.006 unpaired Mann-Whitney t-test) and *CD74-TdT* (Pre: ND, n=3; Post: 0.6 ± 0.2%, n=9, p=0.009 unpaired Mann-Whitney t-test) spinal cord (Figure 2D). In non-immunized *B2m-TdT* adult spinal cord, a small proportion of Olig2+ oligodendroglia were TdT+ (1.3 ± 0.3%, n=4), and these were predominantly mature CC1+ oligodendroglia (88.6 ± 1.7%, n=4) (Figure 2E). In EAE spinal cord, the majority of TdT+Olig2+ oligodendroglia were not immunoreactive to PDGFRa (Figure 2F) in either *B2m-TdT* (93.7 ± 2.4%, n=4) or *CD74-TdT* (83.1 ± 7.1%, n=4) mice, indicating that most of these MHC expressing oligodendroglia had advanced beyond the progenitor stage. Individual animal clinical score (Figure 2G) and region of interest TdT mean fluorescent intensity (Figure 2H), a surrogate for overall inflammation, were both positively correlated with the percentage of TdT+ oligodendroglia, highlighting the close correspondence between activation of MHC pathways in oligodendroglia and disease state. TdT+ oligodendroglia were more abundant in lesion compared to non-lesion areas (Figure 2I) in *B2m-TdT* EAE mice (p=0.039 paired Wilcoxon t-test, n=8) but not *CD74-TdT* EAE mice (p=0.840 paired Wilcoxon t-test, n=10).

**Figure 2.**
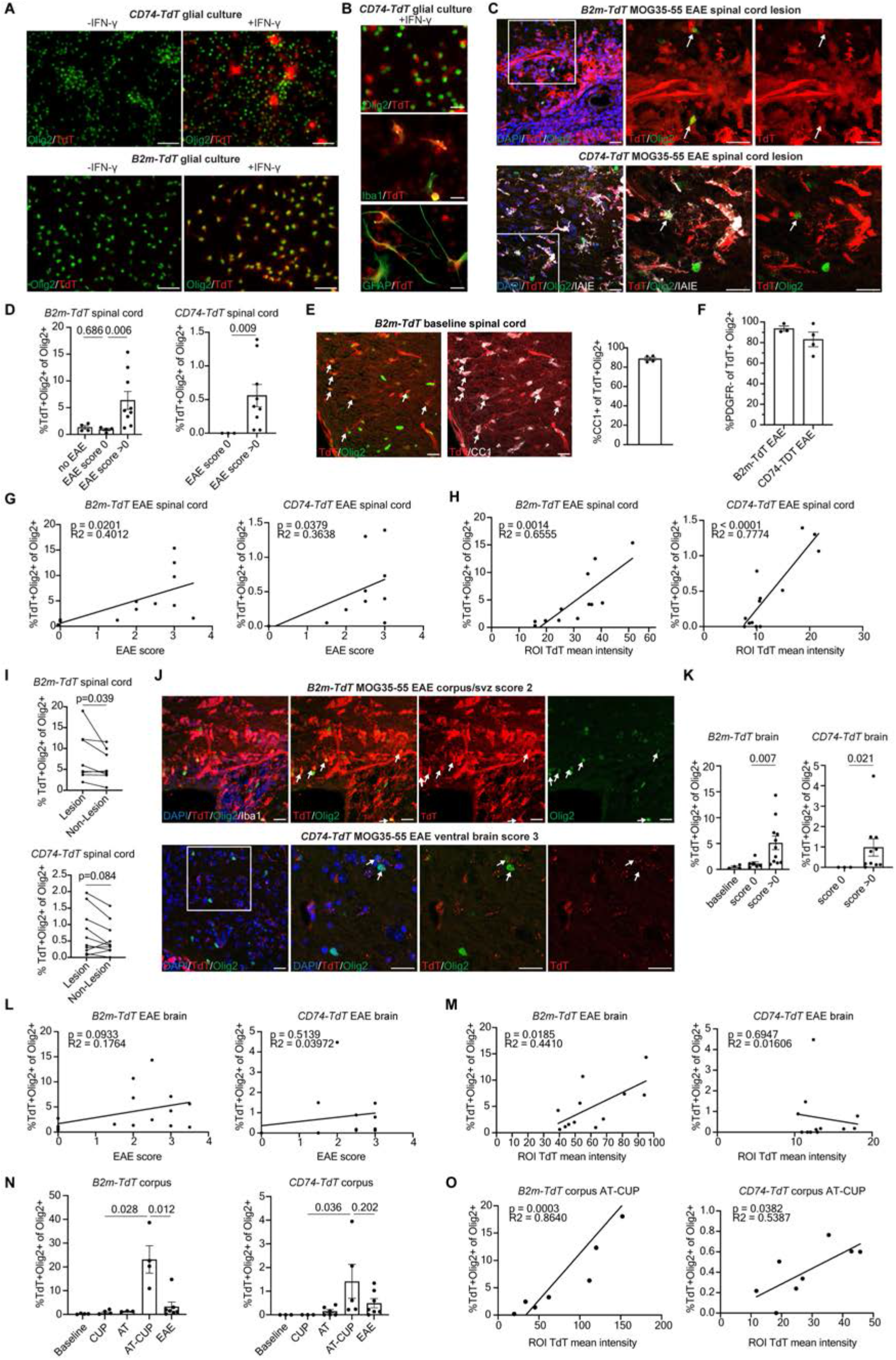
Oligodendroglial MHC reporter expression in CNS inflammatory models. (**A**). Representative images of *CD74-TdT* and *B2m-TdT* postnatal brain immunopanned glial culture stained for TdT and Olig2 with and without IFN-γ treatment. Scale bars, 50μm. (**B**). Representative images of TdT expression in Olig2+ oligodendroglia, GFAP+ astrocytes and Iba1+ microglia in *CD74-TdT* postnatal brain immunopanned glial culture with IFN-γ treatment. Scale bars, 20μm. (**C**). Representative confocal imaging of TdT reporter-positive oligodendroglia in MOG_35-55_ EAE *CD74-TdT* and *B2m-TdT* spinal cord lesions. Endogenous TdT reporter-positive oligodendroglia indicated by arrows. Scale bars, 20μm. (**D**). Quantification of percentage of TdT+Olig2+ oligodendroglia in MOG_35-55_ EAE spinal cord. n = 4 mice no EAE, n = 3-4 mice score 0, n = 9 mice score>0. Unpaired Mann-Whitney t-test. (**E**). Representative confocal imaging of adult *B2m-TdT* spinal cord at baseline with endogenous TdT colocalization with Olig2+CC1+ mature oligodendroglia (arrows). Quantification of percentage of CC1+TdT+Olig2+ oligodendrocytes in *B2m-TdT* no EAE adult spinal cord. n = 4 mice. Scale bars, 20μm. (**F**). Quantification of percentage of PDGFRa-TdT+ oligodendroglia in MOG_35-55_ EAE spinal cord. n = 3-4 mice score>0. (**G**). Linear regression of percentage of TdT+Olig2+ oligodendroglia compared to EAE clinical score in MOG_35-55_ EAE spinal cord. n = 12-13 mice/reporter. (**H**). Linear regression of percentage of TdT+Olig2+ oligodendroglia compared to ROI TdT mean fluorescent intensity in MOG_35-55_ EAE spinal cord. n = 12-13 mice/reporter. (**I**). Quantification of percentage of TdT+Olig2+ oligodendroglia in lesion compared to non-lesion areas in MOG_35-55_ EAE spinal cord. n = 8-10 mice/reporter, paired Wilcoxon t-test. (**J**). Representative confocal imaging of TdT reporter-positive oligodendroglia in MOG_35-55_ EAE *CD74-TdT* and *B2m-TdT* brain. Endogenous TdT reporter-positive oligodendroglia indicated by arrows. Scale bars, 20μm. (**K**). Quantification of percentage of TdT+Olig2+ oligodendroglia in MOG_35-55_ EAE brain. n = 3-6 mice score 0, n = 10-11 mice score>0. Unpaired Mann-Whitney t-test. (**L**). Linear regression of percentage of TdT+Olig2+ oligodendroglia compared to EAE clinical score in MOG_35-55_ EAE brain. n = 13-17 mice/reporter. (**M**). Linear regression of percentage of TdT+Olig2+ oligodendroglia compared to ROI TdT mean fluorescent intensity in MOG_35-55_ EAE brain. n = 12 mice/reporter. (**N**). Quantification of percentage of TdT+Olig2+ oligodendroglia in corpus callosum of baseline adult, cuprizone (CUP), 2D2 Th17 adoptive transfer (AT), cuprizone followed by 2D2 Th17 adoptive transfer (AT-CUP) and MOG EAE (EAE). n = 3-7 mice/group. Unpaired Mann-Whitney t-test. (**O**). Linear regression of percentage of TdT+Olig2+ oligodendroglia compared to ROI TdT mean fluorescent intensity in AT-CUP corpus callosum. n = 8-9 mice/group. Data represented are means ± s.e.m. ROI-region of interest. Figure 2-video 1 Imaris 3D reconstruction of *CD74-TdT* MOG_35-55_ EAE ventral brain confocal Z stack. *CD74-TdT* endogenous expression (red) found within two Olig2-positive (green) oligodendroglia with DAPI counterstain (blue). Figure 2-source data 1 Immunohistochemistry quantification depicted in Figure 2. Sheets are labeled with letter corresponding to data in Figure 2.

TdT+ oligodendroglia were also observed in the EAE brain (Figure 2J and Figure 2-video 1) and were significantly increased in clinical scoring EAE animals compared to pre-clinical score 0 animals in both *B2m-TdT* (Pre: 1.1 ±0.3%, n=6; Post: 5.1 ± 1.3%, n=11, p=0.007 unpaired Mann-Whitney t-test) and *CD74-TdT* mice (Pre: ND, n=3; Post: 1.0 ± 0.4%, n=10, p=0.021 unpaired Mann-Whitney t-test) animals (Figure 2K). However, clinical score was not correlated with the number of TdT+ oligodendroglia (Figure 2L) and region of interest TdT mean fluorescent intensity was only positively correlated with the number of TdT+ oligodendroglia in the brain of *B2m-TdT* EAE mice (Figure 2M).

To determine whether these MHC reporter mice are versatile enough to use with other inflammatory models, we adoptively transferred MOG-reactive Th17 T cells after cuprizone-mediated demyelination (AT-CUP), which results in inflammatory infiltrates and impaired remyelination in the corpus callosum^20^. As seen with MOG_35-55_ EAE, TdT+ oligodendroglia in the corpus callosum were significantly more abundant in both *CD74-TdT* and *B2m-TdT* AT-CUP mice compared to cuprizone alone (CUP) (Figure 2N) and, as in EAE, the percentage of oligodendroglia significantly correlated with region of interest TdT mean fluorescent intensity (Figure 2O).

Isolation of MHC reporter expressing cells reveals distinct transcriptional subpopulations To further define the properties of MHC activated, “immune” oligodendroglia, we performed single-cell RNA sequencing (scRNA-seq) of TdT positive and negative cells sorted from *CD74-TdT* and *B2m-TdT* MOG_35-55_ EAE brains (Figure 3). Single cells were sorted from whole brains based on TdT expression and viability (Figure 3- figure supplement 1A,B) and 10x scRNA-seq was performed on three isolated populations (TdT positive and negative from *CD74-TdT* mice, and TdT positive from *B2m-TdT* mice) with an average sequencing depth of 2,231-2,835 genes per cell. Clustering was validated by assessing known unique transcripts associated with each cell population (Figure 3- figure supplement 1C). Clustering TdT positive and negative populations from *CD74-TdT* mice revealed mostly non-overlapping clusters (Figure 3A). TdT-negative cells were comprised predominantly of T cells, endothelial cells, oligodendroglia and granulocytes, and TdT-positive cells were predominantly microglia, monocytes/macrophages, dendritic cells and B cells (Figure 3B). The TdT-positive sorted sample contained high CD74 and TdT transcript levels (Figure 3C). TdT-positive sorted cells from *B2m-TdT* mice were predominantly monocytes/macrophages, microglia, endothelial cells and T cells (Figure 3D,E), which varied in levels of TdT and B2m transcripts (Figure 3F). The CD74-TdT monocyte/macrophage cluster had distinct separation of TdT-positive and TdT-negative sorted cluster populations (Figure 3A) and differential gene analysis revealed enrichment of MHC class II, M2 activation and macrophage infiltration chemokine, inflammatory signaling receptor, complement and lysosomal protease transcripts in TdT-positive myeloid cells (Figure 3- figure supplement 2A, Figure 3- figure supplement 3A).

**Figure 3.**
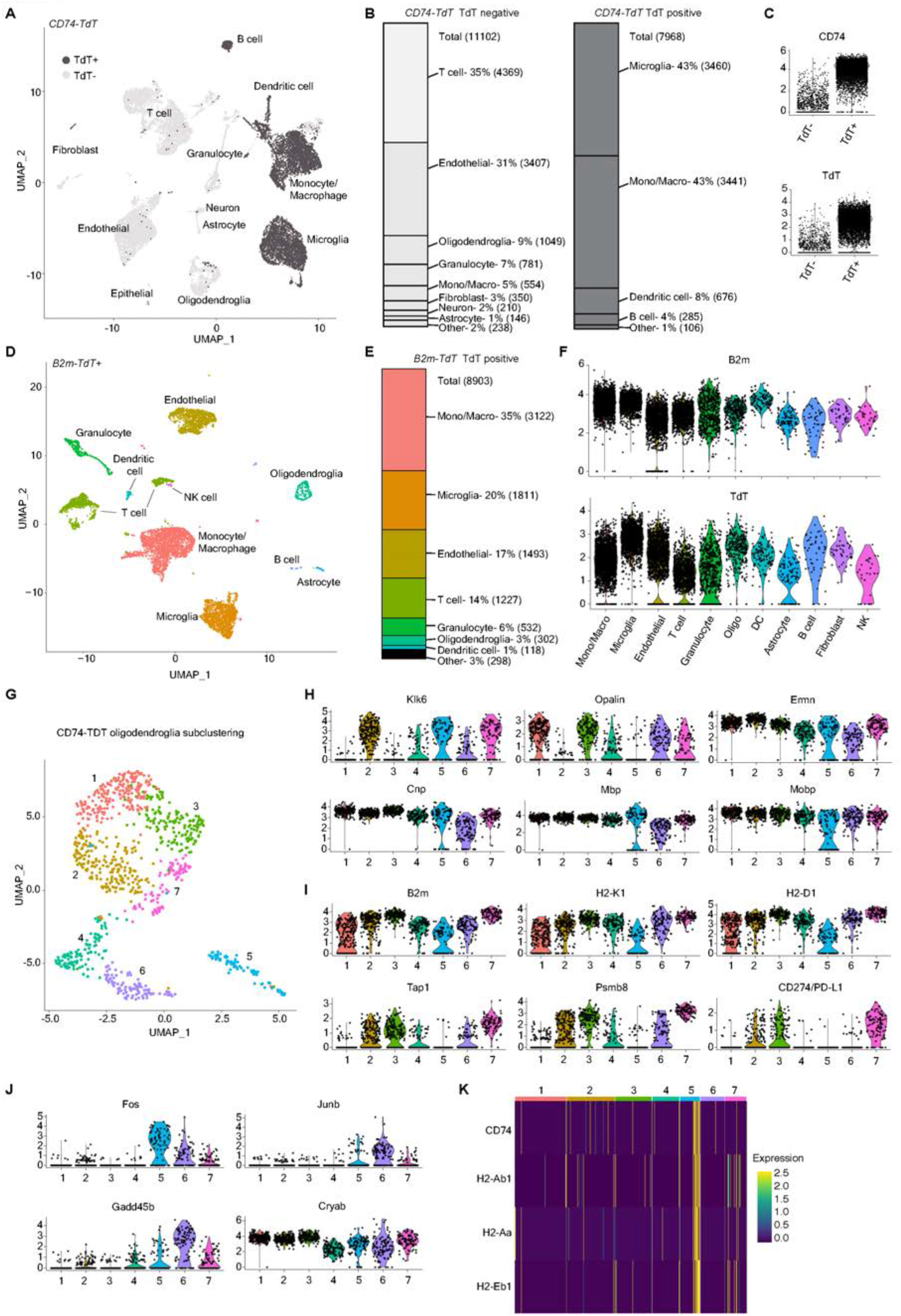
Single-cell RNA sequencing of *CD74-TdT* and *B2m-TdT* MOG_35-55_ EAE brain. (**A**). UMAP *CD74-TdT* MOG_35-55_ EAE brain scRNAseq TdT-positive sorted cells (dark gray) and TdT-negative sorted cells (light gray). (**B**). Cell identity percentages in *CD74-TdT* TdT-positive and TdT-negative sorted cells, total cell count in parentheses. (**C**). Violin plots of log normalized expression levels of CD74 and TdT transcripts in TdT-positive and TdT-negative sorted samples. (**D**). UMAP *B2m-TdT* MOG_35-55_ EAE brain scRNAseq TdT-positive sorted cells. (**E**). Cell identity percentages in *B2m-TdT* TdT-positive sorted cells, cell count in parentheses. (**F**). Violin plots of log normalized expression levels of B2m and TdT transcripts in *B2m-TdT* TdT-positive sorted cell clusters. (**G**). UMAP *CD74-TdT* MOG_35-55_ EAE brain TdT-positive and TdT-negative sorted cells with 7 oligodendroglial subclusters. (**H**). Violin plots of oligodendroglial intermediate (Klk6) and mature (Opalin, Ermn, Cnp, Mbp, Mobp) transcript levels in oligodendroglial subclusters. (**I**). Violin plots of MHC class I (B2m, H2-K1, H2-D1), antigen transport (Tap1), immunoproteasome (Psmb8) and CD274/PD-L1 transcript levels in oligodendroglial subclusters. (**J**). Violin plots of transcripts involved in cell stress (Fos, Junb), growth arrest/apoptosis (Gadd45b) and misfolded protein aggregation (Cryab) in oligodendroglial subclusters. (**K**). Heat map of MHC class II (CD74, H2-Ab1, H2-Aa, H2-Eb1) scaled expression levels in oligodendroglial subclusters. DC-dendritic cells; Mono/Macro-monocyte/macrophage; NK-natural killer. Figure 3- source data 1 Single-cell RNA sequencing data have been deposited in the NCBI’s Expression Omnibus (Edgar et atl., 2002) and are accessible through GEO Series accession number GSE213739 (https://www.ncbi.nlm.nih.gov/geo/query/acc.cgi?acc=GSE213739).

### Oligodendroglia subpopulations in EAE

Subclustering oligodendroglial cells from the *CD74-TdT* EAE brain initially resulted in nine subclusters. On further analysis, several of the clusters with myelin gene transcripts were identified as T cells, myeloid cells and endothelial cells (Figure 3- figure supplement 3B-G, Figure 3- figure supplement 4). The presence of myelin transcripts in these cells may arise through phagocytosis of myelin debris, which has been reported during development^21-22^. After removal of these contaminating clusters, 1,082 oligodendroglia were distributed between seven subclusters (Figure 3G). Differential gene analysis (Figure 3- figure supplement 3H) revealed that clusters 2,4,5,6,7 were enriched for *Klk6* and the lowest levels of mature myelin markers (Figure 3H). *Opalin*, a glycoprotein expressed prior to initiation of myelination^23^ was enriched in clusters 1,3,4,6,7 (Figure 3H). Clusters with the highest MHC class I receptor transcripts (2,3,7) also had the highest levels of immunoproteasome, peptide transport and PD-L1/CD274 transcripts (Figure 3I). It is possible that expression of PD-L1, a MHC class I inhibitory molecule, prevents CD8-mediated killing of these oligodendroglia. Cell stress and growth arrest/apoptosis transcripts were enriched in clusters 5,6,7, which also demonstrated lower levels of chaperone aB-crystallin (Cryab) (Figure 3J), which was previously shown to be upregulated in oligodendrocytes in MS brain^24,25^. The largest group of TdT-positive sorted oligodendroglia (Cluster 5), was notable for expressing the lowest levels of MHC class I receptor transcripts (Figure 3I) and enrichment of MHC class II associated transcripts (Figure 3K).

**Figure 1- figure supplement 5.**
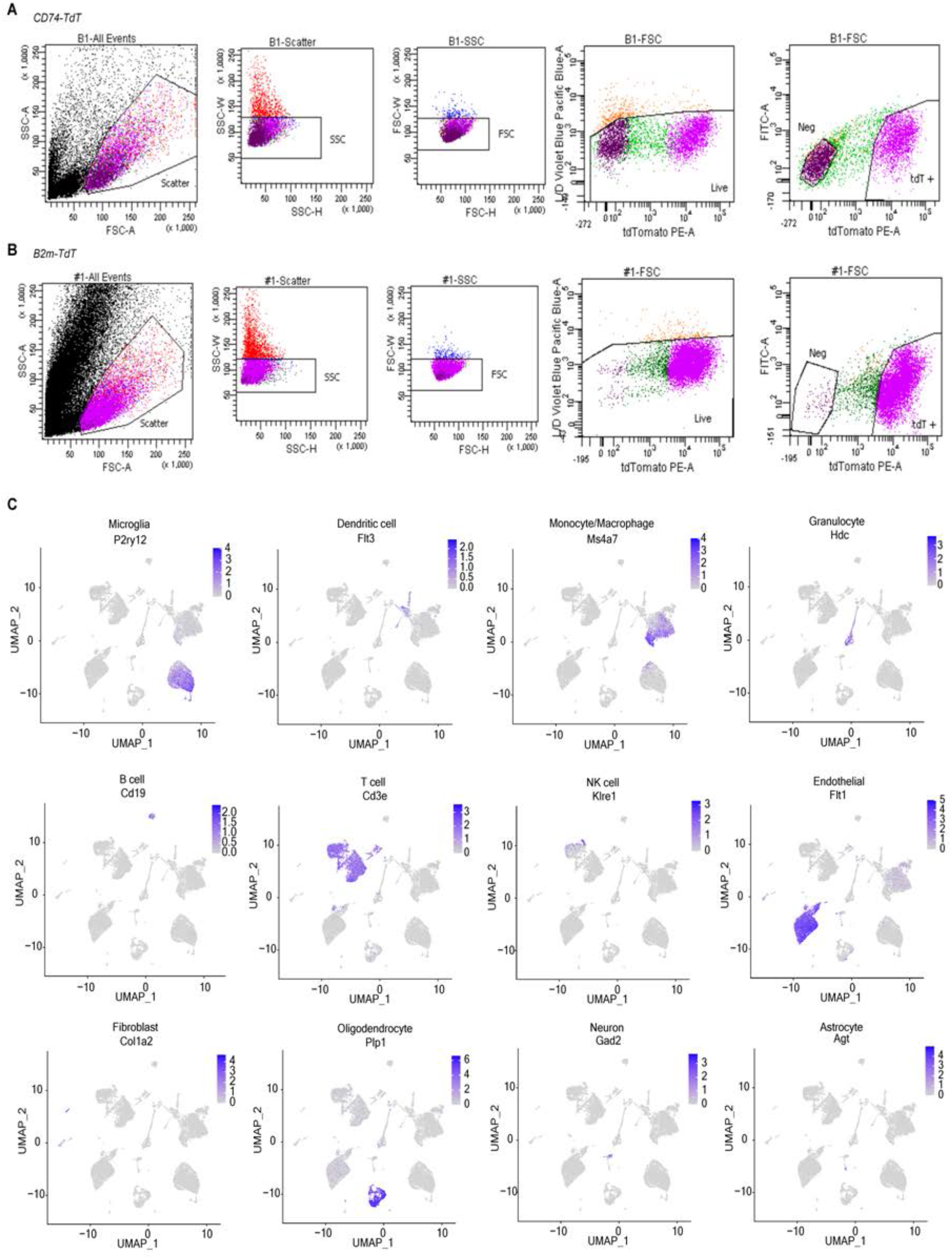
scRNAseq gating strategy and validation of clusters. (**A**). Gating strategy for *CD74-TdT* MOG_35-55_ EAE brains sorted for TdT-positive and TdT-negative cells. (**B**). Gating strategy for *B2m-TdT* MOG_35-55_ EAE brains sorted for TdT-positive and TdT-negative cells. (**C**). Expression level of cell type specific transcripts to confirm cluster identities in *CD74-TdT* MOG_35-55_ EAE.

**Figure 3- figure supplement 2.**
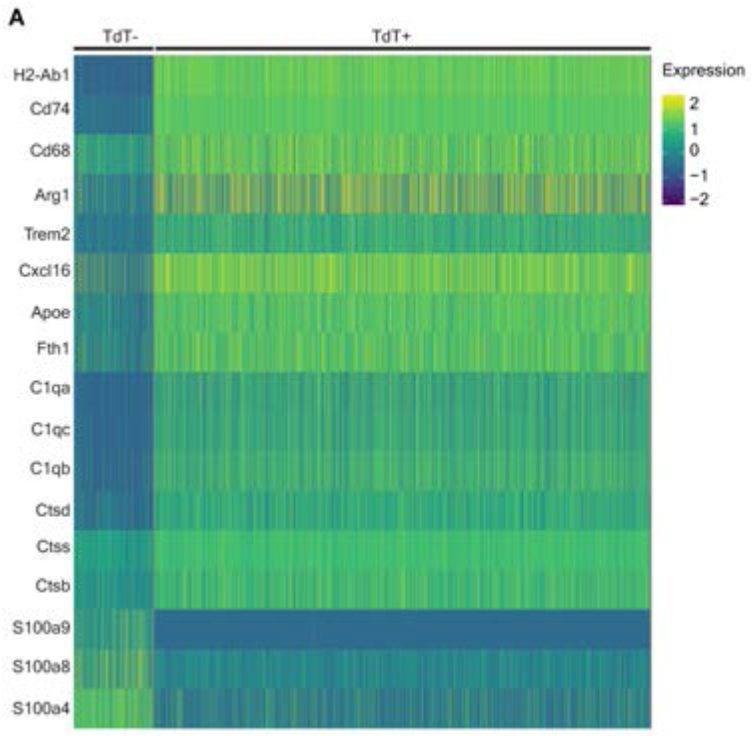
*CD74-TdT* MOG_35-55_ EAE brain scRNAseq monocyte/macrophage cluster differential transcript expression levels. (**A**). Heat map of scaled expression levels in TdT-negative and TdT-positive sorted myeloid/macrophage cluster for MHC class II (H2-Ab1, CD74), M2 activation (CD68, Arg1), inflammatory signaling receptor (Trem2), M2 macrophage infiltration chemokine (Cxcl16), lipid metabolism (Apoe), ferritin light chain (Fth1), complement (C1qa, C1qb, C1qc), lysosomal proteases (Ctsd, Ctss, Ctsb), and calcium sensing transcripts (s100a4, c100a8, s100a9). Figure 3- figure supplement 3 Differential expression transcript analysis of scRNAseq of *CD74-TdT* MOG_35-55_ EAE brain. (**A**). *CD74-TdT* TdT-positive vs TdT-negative sorted myeloid populations differential analysis. Differential expression analysis of contaminating clusters compared to non-contaminating oligodendroglial clusters for myeloid (**B**). T cell (**C**). and endothelial cell (**D**). populations demonstrates enrichment of myeloid, T cell and endothelial cell transcripts, respectively. Differential expression analysis of contaminating cluster compared to cluster of origin that did not cluster with oligodendroglia indicate enrichment of myelin transcripts in contaminating (**E**). myeloid, (**F**). T cells and (**G**). endothelial cells. (**H**). Differential expression analysis of oligodendroglial clusters with contaminating clusters removed with top enriched transcripts for individual oligodendroglial subclusters.

**Figure 3- figure supplement 4.**
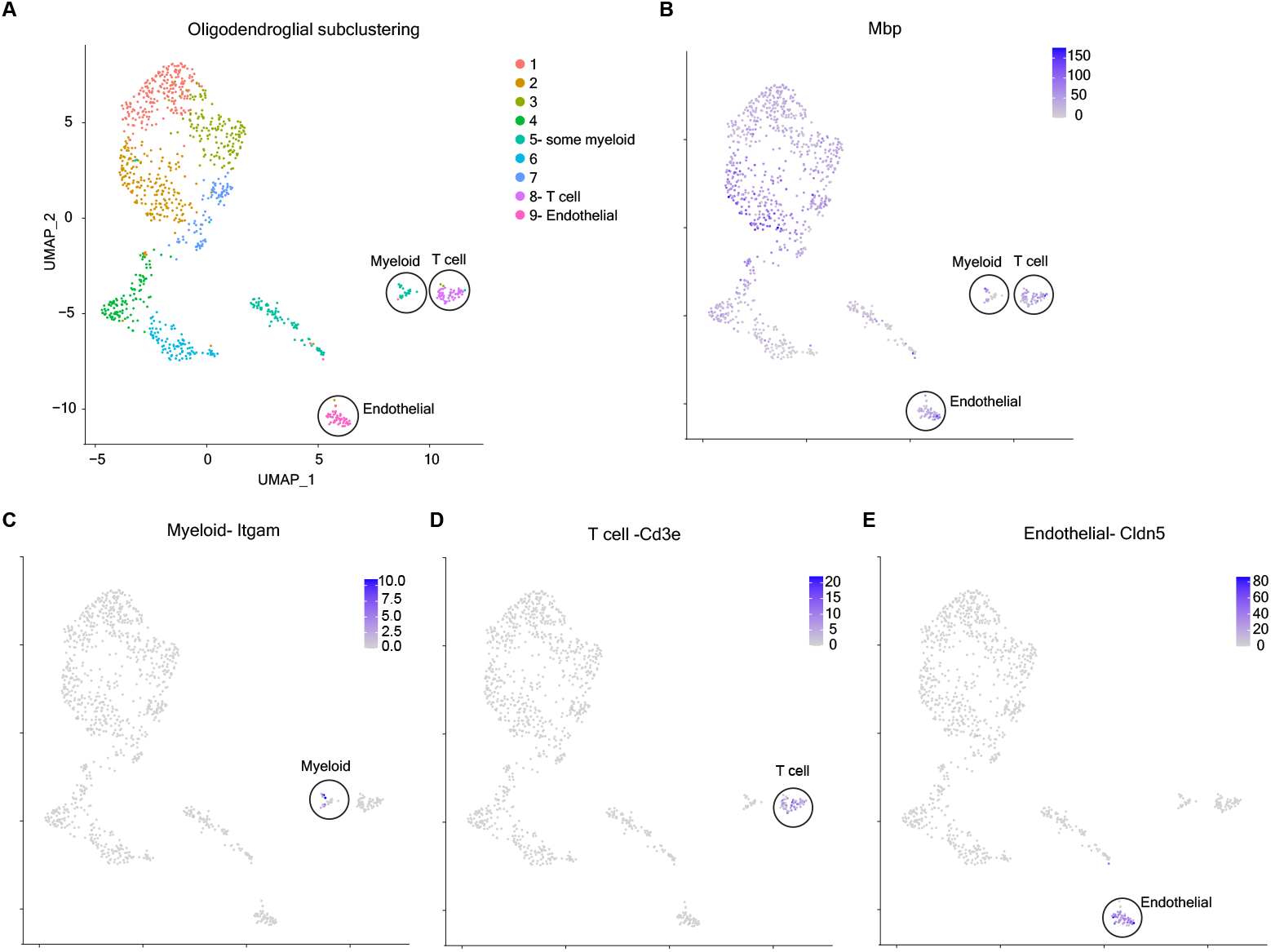
Oligodendroglial subclustering with contaminating clusters in scRNAseq *CD74-TdT* MOG_35-55_ EAE brain TdT-positive and TdT-negative sorted cells. (**A**). UMAP of oligodendroglial subclusters with 9 clusters identified and contaminating clusters outlined. (**B**). UMAP of Mbp transcript levels in oligodendroglial subclusters with myelin transcripts found in outlined contaminating clusters. (**C**). Myeloid contaminating cluster with expression of Itgam transcript. (**D**). T cell contaminating cluster with expression of Cd3e transcript. (**E**). Endothelial contaminating cluster with expression of Cldn5 transcript.

## Discussion

Together, these studies reveal that MHC class I and II reporter mice enable reliable detection of immune oligodendroglia in the CNS. Quantitative analysis from naïve and inflammatory disease model mice reveal that these glial cells represent a small, but diverse population that increase in the inflammatory environment, mirroring single nucleus RNA sequencing results from human MS post-mortem tissues^3-5^. Although not restricted to lesion sites, immune oligodendroglia were higher in areas of enhanced inflammation specific to disease pathology, where they have the potential to influence the behavior of both peripheral and central immune cells and subsequent remyelination. Longitudinal intravital imaging of *CD74-TdT* and *B2m-TdT* mice will help to define the onset and persistence of MHC activation in these cells and determine if they are at higher risk for removal from the CNS through cytotoxic CD8 T cell-induced death. Recent evidence suggests MHC expression may vary with age^26-27^, and our scRNA-seq data revealed that a subset of MHC class I-expressing oligodendroglia express PD-L1/CD274, which may protect these cells from CD8 T cell-mediated death. Conversely, oligodendroglia expressing MHC class II may be more prone to apoptosis or senescence through Gadd45b signaling^28^. Moreover, the frequency of immune oligodendroglia was correlated with disease severity, suggesting that these cells have a relationship with behavioral presentation that could be used to track the progression and treatment of demyelinating disease. The ability to isolate immune oligodendroglia, and other cells with activated MHC pathways from the CNS will help define the phenotypic changes they exhibit in diverse disease and injury contexts, and provide new insight into the complex cellular interactions that modify disease progression and repair.

## Materials and Methods

### Mice

*2D2* (C57BL/6-Tg(Tcra2D2,Tcrb2D2) 1Kuch/J Jax stock #006912) and wild-type C57BL/6 mice were purchased from Jackson Laboratories. Generation of *B2m-TdTomato* and *CD74-TdTomato* reporter lines is described below. All experiments were performed when mice were 8-16 weeks-old and both male and female mice were used. All animal procedures were performed according to protocols approved by the Johns Hopkins Animal Care and Use Committee.

Generation of B2m-TdTomato and CD74-TdTomato reporter lines by Crispr/Cas9 Benchling gRNA design tool was used to select guide RNA (gRNA) sequences to target replacement of the stop codon with reporter repair construct. gRNA sequences were synthesized by Integrated DNA Technologies. Repair reporter constructs were cloned by Genscript and consisted of pUC57 vector with 500bp homology arms of endogenous B2m and CD74 locus flanking P2A-TdTomato-WPRE-pA sequence (repair construct and guide RNA sequence FigS1). Johns Hopkins Transgenic Core Laboratory injected C57BL/6 embryos with gRNA and repair construct (6kb plasmid with 500bp homology arms and 2339 knock-in sequence). Founder pups (62 for *B2m-TdTomato* and 61 for *CD74-TdTomato*) were screened with primers spanning the 5’ homology arm and TdTomato (out-L-F/TdT-R), 3’ homology arm and WPRE sequence (WPRE-F/Out-R-R) and full-length insert 5’ homology arm and 3’ homology arm (In-L-F/In-R-R). For *B2m-TdT*, 5 founder pups demonstrated amplification of all primer sets and 2 of these founder pups had strong TdTomato endogenous fluorescence on peripheral blood flow. Both founder animals were bred to C57BL/6 to generate founder lines and both founders had identical reporter expression in all tissues analyzed. Full length sequencing of insert was unsuccessful due to redundancy in TdTomato sequence. The founder with the strongest PCR amplification band of full-length insert was used for experiments in this study.

For *CD74-TdT*, 1 founder pup demonstrated amplification of all primer sets and had strong TdTomato endogenous fluorescence on peripheral blood flow. This founder animal was bred to C57BL/6 to generate the *CD74-TdT* founder line.

### Oligodendrocyte progenitor culture

Post-natal P6-10 *B2m-TdT* and *CD74-TdT* pups were screened for TdTomato expression with peripheral blood flow. TdT-positive pups were cervically decapitated, forebrains were dissected in HBSS and 3-5 forebrains were pooled and gently chopped several times with a razor. Dissociation was performed with Miltenyi Neural Tissue dissociation kit P (130-092-628) and according to manufacturer’s protocol. After dissociation, cells were resuspended in 0.02% BSA in HBSS. Three 15cm non-coated petri dishes for each dissociation of 3-5 pups were pre-incubated overnight at 4°C the day prior to dissociation. Plate coating: goat anti-rat IgG (Jackson 112-005-167) 1:333, goat anti-mouse IgG (Jackson 115-005-003) 1:333, and BSL1 (Vector L1100) 1:1000 all in Tris-HCl pH 9.5. Prior to dissection and dissociation, secondary antibody plates were washed with HBSS and incubated with primary antibody mouse CD11b (BioRad MCA275G) and rat PDGFRa (BD Pharmigen 558774) at room temperature for at least 2 hours. BSL1 plate was equilibrated at room temperature. After cell dissociation, panning plates were washed with HBSS and cell suspension was added to BSL1 plate (negative selection for endothelial cells) for 10 minutes. Plate was tapped to dislodge loosely bound cells and unbound suspension was transferred to CD11b plate (negative selection for microglia) for 20 minutes and then PDGFRa plate (positive selection for oligodendrocyte progenitors) for 90 minutes. To harvest bound cells, PDGFRa plate was washed with HBSS and 0.0625% trypsin in HBSS was added for 10 minutes at 37°C. FBS was used to dislodge cells and inhibit trypsin and cells were resuspended in OPC proliferation media. Cells were plated at a density of 50k/coverslip with pre-plating to ensure adherence by adding 50k cells in 50ul media to center of coverslip and incubating 37°C for 10 minutes then gently adding 450ul of media to side of well. Coverslips were pre-coated with 10ug/ml PDL in dH_2_O for several hours and washed with dH_2_O prior to cell plating. OPC proliferation media: DMEM high glucose pyruvate (ThermoFisher 11995073), 100U/100ug/ml penicillin/streptomycin, 5ug/ml N-acetyl-L-cysteine, 1xSATO (1ug/ml human apo-transferrin, 1ug/ml BSA, 0.16ug/ml putrescine, 0.6mg/ml progesterone, 0.4ng/ml sodium selenite), 1xB27 (ThermoFisher 17504044), 1xTrace Elements B (Cellgro 99-175-C), 5ug/ml insulin, 10ng/ml d-biotin, 4.2ug/ml forskolin, 20ng/ml PDGFAA (PeproTech 100-13A), 1ng/ml NT-3 (PeproTech 450-03), 10ng/ml CNTF (PeproTech 450-13). Cells were maintained in 37°C incubator with 10% CO_2_ and 50% of media was changed every 48 hours. 24 hours post-plating, 50ng/ml IFN-*γ* was added to media and after 72 hours of IFN-*γ* treatment, coverslips were fixed with 4% PFA for 10 minutes then washed twice in PBS.

### Western blot

Mice were anesthetized with isoflurane and cervically dislocated. Brain, spinal cord, spleen and lymph nodes were dissected and transferred to cold RIPA lysis buffer with complete protease inhibitor cocktail (Millipore Sigma 4693116001) and mortar and pestle was used to grind tissue. Homogenate was centrifuged for 20 minutes 4°C at 12,000rpm, supernatant was collected and protein concentration was measured with BCA assay (Pierce BCA protein assay kit 23225). 20ug of protein samples in Laemmli buffer were reduced with 0.1M DTT and incubated at 100°C for 5 minutes. Samples and Chameleon Duo ladder (Li-Cor 928-60000) were loaded in 4-20% Mini-Protean TGX gel (BioRad) and run at 200V for 45 minutes. Wet transfer was performed onto PVDF membrane at 100V for 60 minutes. Membranes were blocked in Odyssey blocking buffer (Li-Cor 927-60001) for 1 hour at room temperature then incubated in primary antibody in Odyssey blocking buffer with 0.2%Tween-20 overnight at 4°C. Primary antibodies were all used at 1:10,000 concentration: rabbit B2m (Dako A0072), goat TdTomato (MyBiosource MBS448092), rabbit CD74 (Abcam ab245692), and mouse beta-actin (Sigma A2228).

Membranes were washed 3x in TBST (TBS 0.1%Tween-20) for 10 minutes. Membranes were incubated with secondary antibodies in Odyssey blocking buffer with 0.2%Tween-20 for 1 hour at room temperature. Secondary antibodies were all used at 1:10,000 concentration: goat anti-mouse IgG IRDye 680RD (Li-Cor 926-68070), goat anti-rabbit IgG IRDye 800CW (Li-Cor 926-32211), and donkey anti-goat IgG (Li-Cor 926-68074). Membranes were washed 3x in TBST for 10 minutes and imaged on LiCor Odyssey CLX system. Images were exported and band intensity was analyzed in ImageStudioLite software.

### MOG_35-55_ Peptide Experimental Autoimmune Encephalomyelitis (EAE)

MOG_35-55_ peptide was emulsified in 1:1 volume of MOG_35-55_ 2mg/ml in PBS and complete Freund’s adjuvant (8mg/ml mycobacterium tuberculosis in Incomplete Freund’s Adjuvant) with syringes attached to stopcock for 10 minutes. 75ul of emulsion was injected subcutaneously on the left and right lateral abdomen for a total of 150ug MOG_35-55_ peptide. Pertussis toxin 250ng was injected intraperitoneally on day of immunization and 2 days post-immunization (dpi). Clinical scores were monitored daily after 7 dpi. The following criteria for clinical scoring was used from Hooke laboratories: 0.5 mild tail weakness, 1 limp tail, 1.5 limp tail and wobbly walk (no overt leg weakness), 2 limp tail and hindlimb weakness (not dragging), 2.5 limp tail and dragging of one or both hindlimbs but some movement at hindlimb, 3 limp tail and complete hindlimb paralysis (spinning or severe ataxia), 3.5 limp tail and complete hindlimb paralysis and hindlimbs held to one side of body or hindquarters flat, 4 all of 3.5 scoring with partial to complete front leg paralysis and minimal movement around the cage, 5 dead. Mice were sacrificed at 8-17 dpi.

### Flow cytometry

*B2m-TdT* and *CD74-TdT* animals immunized with MOG_35-55_ peptide were sacrificed at EAE clinical score equal to or >3 on 12-15dpi and non-immunized, age-matched *B2m-TdT* and *CD74-TdT* animals were used for baseline tissue. Animals were anesthetized with isoflurane and intracardially perfused with cold HBSS. Spleens were dissected and dissociated by mashing over 100um filter and red blood cells were lysed using RBC lysis buffer (BioLegend 420301). Brain and spinal cords were dissected, cut with razor sagittally for two dissociations per animal, lightly chopped several times with fine razor and transferred to pre-equilibrated enzyme solution at 37°C. Two separate dissociations with papain/DNase and collagenase/DNase were performed for each sagittal half of the brain and spinal cord. Concentrations of enzymes: papain 20U/ml (Worthington LS003119), collagenase IV 1mg/ml (Worthington LS004188) and DNase 100U/ml (Worthington LS002007). Papain buffer: EBSS (Sigma E7510), 22.5mM D-glucose, 0.5mM EDTA, 2.2g/L NaHCO_3_, 5.5mM L-cysteine pH 7.4. Collagenase buffer: EBSS, 22.5mM D-glucose, 2.2g/L NaHCO_3_, 3mM CaCl_2_ pH 7.4. Tissue dissociations were incubated at 37°C for 10 minutes followed by mechanical dissociation 10x with 1ml pipette tip (first time with tip cut to larger bore) and repeated for a total of three incubations/triturations. Cell dissociations were filtered over a 70um filter, washed with PBS and myelin was removed with debris removal solution (Miltenyi 130-109-398) according to manufacturer’s protocol. Lower cell suspension layer was washed with PBS, filtered over FACS tube with 35um filter, incubated for 15 minutes with Fc block (BioLegend 156604) and live/dead Aqua (ThermoFisher L34966) for spleen samples, washed with FACs buffer and incubated with cell surface antibodies at concentration of 1:100 in FACs buffer for 30 minutes. Cell surface antibodies: CD45 APCFire 750 (BioLegend 103211), Ly6G bv650 (BioLegend 127641), CD19 PacBlue (BioLegend 115523), CD11b PerCPCy5.5 (BioLegend 101228), CD11c PECy7 (BioLegend 117318), Clec12a APC (BioLegend 143406), CD86 bv421 (BioLegend 105035), CD40 FITC (BioLegend 124608), CD206 bv711 (BioLegend 141727), IA/IE bv785 (BioLegend 107645), CD3 PerCPCy5.5 (BioLegend 317336), CD8 PECy7 (BioLegend 100722), CD4 APC (BD 553051). For B2m staining, rabbit B2m 1:10 (Agilent A0072) primary was added to surface stain, and 3 washes were performed prior to secondary antibody staining with anti-rabbit bv421 1:500 (BioLegend 406410) with 3 additional washes after secondary antibody. For CD74 staining after surface stain, cells were fixed with IC fix (eBioscience88-8824-00) for 30 minutes followed by incubation with CD74 FITC 1:100 (BD 561941) in permeabilization buffer (eBioscience88-8824-00) for 30min according to manufacturer’s protocol. Cells were washed in FACS buffer and ran on Cytek Aurora 4 laser (VBYR) flow cytometer in Johns Hopkins Ross Flow Cytometry Core. Compensation was performed with single stained UltraComp eBeads (ThermoScientific 01-3333-42), ArC amine reactive beads for viability dyes (LifeTechnologies A10346) and unstained wild-type dissociated spleen, brain and spinal cord for TdTomato compensation. FlowJo software was used for analysis and gating. Spleen was gated on singlet, cell, viable, CD45+, Ly6G-then further subgated on CD19+ (B cells), CD11b+ (monocytes), CD11b-CD11c+ (dendritic cells). For brain and spinal cord EAE cells were gated on singlets, cell, CD45+CD11b+ and then further subgated on Clec12a+ vs Clec12a-. Both Clec12a+ (infiltrating myeloid) and Clec12a-(microglia) were gated on costimulatory molecule CD40 and CD86 expression. Collagenase and papain dissociations were compared without any notable differences and dissociations with the highest yield of cells of interest were presented (collagenase for *CD74-TdT* infiltrating myeloid cells and microglia and papain for *B2m-TdT* microglia).

### Single-cell RNA sequencing

Three MOG_35-55_ peptide immunized *B2m-TdT* male 16-week-old animals were sacrificed at 13dpi with clinical scores of 2.5, 4 and 4. Four MOG_35-55_ peptide immunized *CD74-TdT* female 15-week-old animals were sacrificed at 14dpi with clinical scores of 2.5, 3, 3.5, 3.5. Animals were anesthetized with isoflurane and intracardially perfused with cold HBSS without cations with 5ug/ml actinomycin (Sigma A1410) and 10uM triptolide (Sigma T3652). Brains and spinal cords were dissected and placed in 6 well plate with cold dissection buffer (HBSS no cations with 5ug/ml actinomycin, 10uM triptolide, and 27ug/ml anisomycin (Sigma A9789)) protected from light. Brains and spinal cords were lightly chopped several times with fine razor and transferred to pre-equilibrated at 37°C dissociation buffer with enzymes (papain 20U/ml and DNase 100U/ml with 5ug/ml actinomycin, 10uM triptolide, and 27ug/ml anisomycin in EBSS, 22.5mM D-glucose, 0.5mM EDTA, 2.2g/L NaHCO_3_, 5.5mM L-cysteine pH 7.4). Tissue dissociations were incubated at 37°C for 10 minutes followed by mechanical dissociation 10x with 1ml pipette tip (first time with tip cut to larger bore) and repeated for a total of three incubations/triturations. Cell dissociations were filtered over a 70um filter, washed with PBS and myelin was removed with two rounds of debris removal solution (Miltenyi 130-109-398) according to manufacturer’s protocol. Cell pellet was resuspended in PBS and filtered over FACS tube with 35um filter and incubated with Zombie live/dead Violet (Biolegend 423114) for 15 minutes in cold PBS at 4°C, washed in cold PBS and resuspended in 0.5% BSA 1mM EDTA PBS at concentration of 1×106 cells/ml in cold PBS on ice for single cell sorting. For *CD74-TdT* samples, two animals were pooled for two sorts. For *B2m-TdT* samples, all three samples were combined then divided into two tubes and only one tube was sorted. Only brain samples were sorted due to the level of myelin debris in spinal cord samples. Brain samples were sorted on Aria Ilu 3 laser (VBR) in Johns Hopkins Ross Flow Cytometry core and gated on scatter and viability. TdT-positive and TdT-negative populations were collected in 0.1% BSA PBS on ice. Samples were sorted for a limit of 2 hours before submission to Johns Hopkins sequencing core. Cell viability and concentration was checked on a Countess II automated hemocytometer using Trypan Blue (Life Technologies) exclusion. Cell volumes calculated to capture 10,000 cells were loaded into a Chromium Next GEM Chip G and GEMs (Gel Bead-in-emulsion) were generated using a Chromium Controller using Single Cell 3’ v3 chemistry (10X Genomics). Barcoded single cell libraries were then generated according to manufacturer recommendations. Libraries were sequenced on a NovoSeq 6000 (Illumina). Sequencing data was aligned to the mouse genome^29^ modified to include the p2A-tdTomato sequence (named TDT in the reference) using cellranger (v6.1.0, 10X Genomics). Filtered features from cellranger were imported into R (v4.1.2) using Seurat^30^ (v4.1.0). Additional per cell metrics were assessed including gene count, UMI count, and ratio of reads mapping to mitochondrial genes. Cells were excluded for gene and UMI counts using adaptive thresholds (+/-3 median absolute deviations) with scuttle^31^ (v1.4.0) or for mitochondrial gene ratios greater than 10%. The CD74-tdT negative and CD74-tdT positive datasets were then merged into a single object prior to data normalization. For clustering purposes, UMI counts were normalized with a regularized negative binomial regression via the SCTransform function in Seurat^32^ using the glmGamPoi method (v1.6.0)^33^. This was followed by principle component analysis (PCA) dimensionality reduction followed by Uniform Manifold Approximation and Projection (UMAP) dimensionality reduction on the first 40 principle components. Clustering was performed by creating a shared nearest-neighbor graph using the first 40 principle components followed by Louvain clustering. Cluster annotation was done in an unbiased fashion with SingleR (v1.8.1)^34^ against the MouseRNASeqData set of 358 mouse bulk RNA-seq samples from sorted populations^35^ available in the celldex package (v1.4.0). Unbiased annotations were then checked manually by comparing against well characterized reference genes. For purposes of expression visualization (violin plots, feature plots), UMI counts were log normalized with Seurat. Subclustering analysis of oligodendrocyte lineage cells was performed by subsetting the oligodendrocytes from the larger merged CD74-tdT dataset and repeating the above steps on just the cells already identified as oligodendrocytes. The authors thank Linsa Orzolek and Tyler Creamer of the JHMI Single Cell & Transcriptomics Core for assistance generating and sequencing scRNAseq libraries. The authors alos thank JHU Ross Flow Cytometry Core for assistance with FACs and for the use of 4 laser Cytek Aurora. The Core Aurora is funded by NIH grant S10OD026859.

### Cuprizone Th17 Adoptive Transfer

Mice were fed 0.2% cuprizone (bis(cyclo-hexanone) oxaldihydrazone (Sigma-Aldrich)) mixed with powdered, irradiated 18% protein rodent diet (Teklad Global) for 3.5 weeks, and chow was replaced every 2-3 days. CD4+ T cells were isolated from the spleens and draining lymph nodes of 2D2 mice using the CD4+ isolation kit (BioLegend) and co-cultured with irradiated wild-type splenocytes at a ratio of 1:5. To polarize cells to a Th17 profile, 2.5 ug/mL anti-CD3 (BioLegend), 20 ug/mL anti-IL-4 (BioLegend), 20 ug/mL anti-IFN-*γ* (BioLegend), 30 ng/mLIL-6 (PeproTech), and 3 ng/mL TGFb (Thermo Fisher) were added to the medium IMDM (Thermo Fisher), fetal bovine serum (Gemini Bio-Products), penicillin and streptomycin (Quality Biologicals), 2-mercaptoethanol (Gibco), Glutamax (Thermo Fisher), and sodium pyruvate (Millipore Sigma)). After 72 hours, cells were transferred to a new plate with fresh medium and IL-23 (R&D Systems). After 48 hours rest, cells were restimulated on plates coated with anti-CD3 and anti-CD28 (BioLegend). After 48 hours of restimulation, cells were collected and resuspended in PBS. Mice fed 0.2% cuprizone for 3.5 weeks (AT-CUP) followed by 3 days of cessation of cuprizone and no cuprizone treatment (AT only) mice were injected intraperitoneally with 10×10^6^ Th17 cells. Clinical score was monitored daily as described above starting at 7 days post adoptive transfer and mice were sacrificed 7-17 days after adoptive transfer. Cuprizone without adoptive transfer was performed in parallel, and mice were sacrificed 7 days after cessation of cuprizone diet.

### Immunohistochemistry

Mice were anesthetized with isoflurane and intracardially perfused with PBS followed by 4% PFA. Brain and spinal cord were dissected and post-fixed in 4% PFA for 24 hours then transferred to 30% sucrose for 2-3 days. Tissue was embedded in OCT (Tissue-Tek) and flash frozen on dry ice bath. Tissue was cut on Leica CM1850 cryostat and spinal cord 20um sections were mounted onto slides and brain 30um sections were mounted onto slides or collected in cryoprotectant (30% sucrose, 30% ethylene glycol, 10mg/ml polyvinylpyrrolidone in 0.1M PB buffer pH 7.4-10.9mg/ml Na_2_PO_4_, 3.2mg/ml NaH_2_PO_4_ in dH_2_O). Floating sections were stored at −20°C and mounted sections were stored at −80°C. Floating sections were mounted onto slides prior to staining and allowed to dry for 10 minutes. Coverslip staining was performed in 24 well plate. Slides were washed with PBST (PBS Triton 0.5%) for 5 minutes, outlined with PAP pen (ThermoFisher 5027627) and blocked with 5% normal goat serum (Vector S-1000) or 5% normal donkey serum (Jackson 017-000-121) in PBST for 1 hour. Slides were incubated in primary antibody in 5% normal goat or normal donkey serum in PBST overnight at 4°C. Primary antibodies: rabbit Olig2 1:1000 (Millipore Sigma AB9610), mouse Olig2 1:1000 (Millipore Sigma MABN50), mouse CC1/APC 1:500 (Millipore Sigma OP80), rabbit PDGFRa 1:500 (Cell Signaling 3174), rabbit Iba1 1:1000 (Wako 019-19741) goat Iba1 1:500 (Novus NB100-1028), goat TdTomato 1:2000 (MyBiosource MBS448092), rat IA/IE 1:100 (BD Pharmingen 556999), rabbit CD74 1:100 (Abcam ab245692), rabbit CD31 1:100 (Abcam ab28364), hamster CD11c 1:100 (ThermoFisher 14-0114-82), TMEM119 (Abcam ab209064), rabbit NeuN 1:1000 (Millipore Sigma ABN78), mouse NeuN 1:500 (Millipore Sigma MAB377), mouse Parvalbumin 1:1000 (Millipore Sigma P3088), rabbit GFAP 1:1000 (Agilent Z0334). Slides were washed 3x PBST then incubated with secondary antibody 1:500 in in 5% normal goat or normal donkey serum in PBST for 1 hour at room temperature. Secondary antibodies: goat anti-rabbit IgG 488 (ThermoScientific A11008) and 647 (ThermoScientific A21244), goat anti-rat IgG 488 (ThermoScientific A11006) and 647 (ThermoScientific A21247), goat anti-hamster IgG 488 (ThermoScientific A21110), goat anti-mouse IgG 488 (ThermoScientific A11001) and 647 (ThermoScientific A21235), donkey anti-goat IgG 544 (Abcam ab150134) and 647 (ThermoScientific A21447). Slides were washed 1x in PBST for 5 minutes, 2x PBS for 5 minutes, then coverslipped with Prolong Gold Antifade with DAPI (Life Technologies P36931).

### Microscopy

Epifluorescence images were taken on Zeiss Axio Observer Z1 with 20x objective. Tiled images were collected with multi-focal Z support points and stitched in Zeiss Zen Blue software.Confocal Z-stack images were taken on Zeiss 880 confocal with 40x objective in Johns Hopkins NINDS Multiphoton Imaging Core NS050274. Maximum intensity projections of Z-stacks were created in Zeiss Zen Black software. Imaris software was used for 3D video rendering of Z stacks.

Oligodendroglial TdT quantification of baseline, MOG_35-55_ peptide EAE, cuprizone, adoptive transfer, and adoptive transfer cuprizone animals

Four brain and lumbar spinal cord sections were analyzed per animal. Z-stack images were taken at 20x magnification at the following ROIs: brain-midline corpus callosum and bilaterally at the corpus horn, subventricular zone, and ventral brain; spinal cord-central canal and bilateral dorsal horn and ventral white matter. The total number of Olig2+ cells and Olig2+TdT+ cells in each image were manually counted for each image using the events feature in Zen Blue software. Regional means were calculated and total mean represented was mean of combined regional means. Mean TdT fluorescent intensity was calculated on non-thresholded 8-bit images using ImageJ. The mean TdT fluorescent intensity represented the mean of regional means for an individual animal. To compare lesion to non-lesion images, the presence of DAPI hypercellularity was used to categorize each image from animals with clinical score >0 as lesion or non-lesion. The mean percentage of Olig2+TdT+ of lesion and non-lesion images across all spinal cord images for each animal was calculated. For animals with clinical score >0 the mean percentage of PDGFRa+Olig2+TdT+ was quantified across all regions. For cuprizone, adoptive transfer and cuprizone adoptive transfer animals, four brain sections were analyzed per animal. Quantification was performed as described above for mid-corpus and bilateral corpus horn regions.

### Statistical analysis

GraphPad prism software was used for statistical analysis. For flow cytometry data unpaired t-tests were used to compare groups. For EAE lesion analysis unpaired Mann-Whitney t-test was used to compare groups given the variation in clinical scores combined into one group. For comparison of same animal lesion to non-lesion a paired Wilcoxon t-test was used. Simple linear regression was used for comparison of Olig2+TdT+ cells to EAE clinical score and mean TdT gray value.

## References

1. Falcão, A.M., van Bruggen, D., Marques, S., Meijer, M., Jäkel, S., Agirre, E., Samudyata, Floriddia, E.M., Vanichkina, D.P., Ffrench-Constant, C., Williams, A., Guerreiro-Cacais, A.O., Castelo-Branco, G., 2018. Disease-specific oligodendrocyte lineage cells arise in multiple sclerosis. Nat. Med. 24, 1837–1844. https://doi.org/10.1038/s41591-018-0236-y

2. Meijer, M., Agirre, E., Kabbe, M., van Tuijn, C.A., Heskol, A., Zheng, C., Mendanha Falcão, A., Bartosovic, M., Kirby, L., Calini, D., Johnson, M.R., Corces, M.R., Montine, T.J., Chen, X., Chang, H.Y., Malhotra, D., Castelo-Branco, G., 2022. Epigenomic priming of immune genes implicates oligodendroglia in multiple sclerosis susceptibility. Neuron 110, 1193–1210.e13. https://doi.org/10.1016/j.neuron.2021.12.034

3. Schirmer, L., Velmeshev, D., Holmqvist, S., Kaufmann, M., Werneburg, S., Jung, D., Vistnes, S., Stockley, J.H., Young, A., Steindel, M., Tung, B., Goyal, N., Bhaduri, A., Mayer, S., Engler, J.B., Bayraktar, O.A., Franklin, R.J.M., Haeussler, M., Reynolds, R., Schafer, D.P., Friese, M.A., Shiow, L.R., Kriegstein, A.R., Rowitch, D.H., 2019. Neuronal vulnerability and multilineage diversity in multiple sclerosis. Nature 573, 75–82. https://doi.org/10.1038/s41586-019-1404-z

4. Jäkel, S., Agirre, E., Mendanha Falcão, A., van Bruggen, D., Lee, K.W., Knuesel, I., Malhotra, D., Ffrench-Constant, C., Williams, A., Castelo-Branco, G., 2019. Altered human oligodendrocyte heterogeneity in multiple sclerosis. Nature 566, 543–547. https://doi.org/10.1038/s41586-019-0903-2

5. Absinta, M., Maric, D., Gharagozloo, M., Garton, T., Smith, M.D., Jin, J., Fitzgerald, K.C., Song, A., Liu, P., Lin, J.-P., Wu, T., Johnson, K.R., McGavern, D.B., Schafer, D.P., Calabresi, P.A., Reich, D.S., 2021. A lymphocyte-microglia-astrocyte axis in chronic active multiple sclerosis. Nature 597, 709–714. https://doi.org/10.1038/s41586-021-03892-7

6. Lau, S.-F., Cao, H., Fu, A.K.Y., Ip, N.Y., 2020. Single-nucleus transcriptome analysis reveals dysregulation of angiogenic endothelial cells and neuroprotective glia in Alzheimer’s disease. Proc. Natl. Acad. Sci. U. S. A. 117, 25800–25809. https://doi.org/10.1073/pnas.2008762117

7. Pan, R., Zhang, Q., Anthony, S.M., Zhou, Y., Zou, X., Cassell, M., Perlman, S., 2020. Oligodendrocytes that survive acute coronavirus infection induce prolonged inflammatory responses in the CNS. Proc. Natl. Acad. Sci. U. S. A. 117, 15902–15910. https://doi.org/10.1073/pnas.2003432117

8. Malone, K.E., Stohlman, S.A., Ramakrishna, C., Macklin, W., Bergmann, C.C., 2008. Induction of class I antigen processing components in oligodendroglia and microglia during viral encephalomyelitis. Glia 56, 426–435. https://doi.org/10.1002/glia.20625

9. Phares, T.W., Ramakrishna, C., Parra, G.I., Epstein, A., Chen, L., Atkinson, R., Stohlman, S.A., Bergmann, C.C., 2009. Target-dependent B7-H1 regulation contributes to clearance of central nervous system infection and dampens morbidity. J. Immunol. 182, 5430–5438. https://doi.org/10.4049/jimmunol.0803557

10. Harrington, E.P., Bergles, D.E., Calabresi, P.A., 2020. Immune cell modulation of oligodendrocyte lineage cells. Neurosci. Lett. 715, 134601. https://doi.org/10.1016/j.neulet.2019.134601

11. Kirby, L., Jin, J., Cardona, J.G., Smith, M.D., Martin, K.A., Wang, J., Strasburger, H., Herbst, L., Alexis, M., Karnell, J., Davidson, T., Dutta, R., Goverman, J., Bergles, D., Calabresi, P.A., 2019. Oligodendrocyte precursor cells present antigen and are cytotoxic targets in inflammatory demyelination. Nat. Commun. 10, 3887. https://doi.org/10.1038/s41467-019-11638-3

12. Lubetzki, C., Zalc, B., Williams, A., Stadelmann, C., Stankoff, B., 2020. Remyelination in multiple sclerosis: from basic science to clinical translation. Lancet Neurol. 19, 678–688. https://doi.org/10.1016/S1474-4422(20)30140-X

13. Mahad, D.H., Trapp, B.D., Lassmann, H., 2015. Pathological mechanisms in progressive multiple sclerosis. Lancet Neurol. 14, 183–193. https://doi.org/10.1016/S1474-4422(14)70256-X

14. Lassmann, H., van Horssen, J., Mahad, D., 2012. Progressive multiple sclerosis: pathology and pathogenesis. Nat. Rev. Neurol. 8, 647–656. https://doi.org/10.1038/nrneurol.2012.168

15. Su, H., Na, N., Zhang, X., Zhao, Y., 2017. The biological function and significance of CD74 in immune diseases. Inflamm. Res. 66, 209–216. https://doi.org/10.1007/s00011-016-0995-1

16. Schröder, B., 2016. The multifaceted roles of the invariant chain CD74--More than just a chaperone. Biochim. Biophys. Acta 1863, 1269–1281. https://doi.org/10.1016/j.bbamcr.2016.03.026

17. Jurga, A.M., Paleczna, M., Kuter, K.Z., 2020. Overview of General and Discriminating Markers of Differential Microglia Phenotypes. Front. Cell. Neurosci. 14, 198. https://doi.org/10.3389/fncel.2020.00198

18. Fancy, S.P.J., Harrington, E.P., Yuen, T.J., Silbereis, J.C., Zhao, C., Baranzini, S.E., Bruce, C.C., Otero, J.J., Huang, E.J., Nusse, R., Franklin, R.J.M., Rowitch, D.H., 2011. Axin2 as regulatory and therapeutic target in newborn brain injury and remyelination. Nat. Neurosci. 14, 1009–1016. https://doi.org/10.1038/nn.2855

19. Shrestha, B., Jiang, X., Ge, S., Paul, D., Chianchiano, P., Pachter, J.S., 2017. Spatiotemporal resolution of spinal meningeal and parenchymal inflammation during experimental autoimmune encephalomyelitis. Neurobiol. Dis. 108, 159–172. https://doi.org/10.1016/j.nbd.2017.08.010

20. Baxi, E.G., DeBruin, J., Tosi, D.M., Grishkan, I.V., Smith, M.D., Kirby, L.A., Strasburger, H.J., Fairchild, A.N., Calabresi, P.A., Gocke, A.R., 2015. Transfer of myelin-reactive th17 cells impairs endogenous remyelination in the central nervous system of cuprizone-fed mice. J. Neurosci. 35, 8626–8639. https://doi.org/10.1523/JNEUROSCI.3817-14.2015

21. Irfan, M., Evonuk, K.S., DeSilva, T.M., 2022. Microglia phagocytose oligodendrocyte progenitor cells and synapses during early postnatal development: implications for white versus gray matter maturation. FEBS J. 289, 2110–2127. https://doi.org/10.1111/febs.16190

22. Nemes-Baran, A.D., White, D.R., DeSilva, T.M., 2020. Fractalkine-Dependent Microglial Pruning of Viable Oligodendrocyte Progenitor Cells Regulates Myelination. Cell Rep. 32, 108047. https://doi.org/10.1016/j.celrep.2020.108047

23. Golan, N., Adamsky, K., Kartvelishvily, E., Brockschnieder, D., Möbius, W., Spiegel, I., Roth, A.D., Thomson, C.E., Rechavi, G., Peles, E., 2008. Identification of Tmem10/Opalin as an oligodendrocyte enriched gene using expression profiling combined with genetic cell ablation. Glia 56, 1176–1186. https://doi.org/10.1002/glia.20688

24. van Noort, J.M., van Sechel, A.C., Bajramovic, J.J., el Ouagmiri, M., Polman, C.H., Lassmann, H., Ravid, R., 1995. The small heat-shock protein alpha B-crystallin as candidate autoantigen in multiple sclerosis. Nature 375, 798–801. https://doi.org/10.1038/375798a0

25. Bsibsi, M., Peferoen, L.A.N., Holtman, I.R., Nacken, P.J., Gerritsen, W.H., Witte, M.E., van Horssen, J., Eggen, B.J.L., van der Valk, P., Amor, S., van Noort, J.M., 2014. Demyelination during multiple sclerosis is associated with combined activation of microglia/macrophages by IFN-γ and alpha B-crystallin. Acta Neuropathol. 128, 215–229. https://doi.org/10.1007/s00401-014-1317-8

26. de la Fuente, A.G., Queiroz, R.M.L., Ghosh, T., McMurran, C.E., Cubillos, J.F., Bergles, D.E., Fitzgerald, D.C., Jones, C.A., Lilley, K.S., Glover, C.P., Franklin, R.J.M., 2020. Changes in the Oligodendrocyte Progenitor Cell Proteome with Ageing. Mol. Cell. Proteomics 19, 1281–1302. https://doi.org/10.1074/mcp.RA120.002102

27. Mishra, A., Shang, Y., Wang, Y., Bacon, E.R., Yin, F., Brinton, R.D., 2020. Dynamic Neuroimmune Profile during Mid-life Aging in the Female Brain and Implications for Alzheimer Risk. iScience 23, 101829. https://doi.org/10.1016/j.isci.2020.101829

28. Zaidi, M.R., Liebermann, D.A., 2022. Gadd45 in Senescence. Adv. Exp. Med. Biol. 1360, 109–116. https://doi.org/10.1007/978-3-030-94804-7_8

29. Howe K.L., Achuthan P., Allen J., Allen J., Alvarez-Jarreta J., Amode M.R., Armean I.M., Azov A.G,. Bennett R., Bhai J., Billis K., Boddu S., Charkhchi M., Cummins C., Da Rin Fioretto L., Davidson C., Dodiya K., El Houdaigui B., Fatima R., Gall A., Garcia Giron C., Grego T., Guijarro-Clarke C., Haggerty L., Hemrom A., Hourlier T., Izuogu O.G., Juettemann T., Kaikala V., Kay M., Lavidas I., Le T., Lemos D., Gonzalez Martinez J., Marugán J.C., Maurel T., McMahon A.C., Mohanan S., Moore B., Muffato M., Oheh D.N., Paraschas D., Parker A., Parton A., Prosovetskaia I., Sakthivel M.P., Salam A.I.A., Schmitt B.M., Schuilenburg H, Sheppard D, Steed E, Szpak M, Szuba M, Taylor K, Thormann A, Threadgold G., Walts B., Winterbottom A., Chakiachvili M., Chaubal A., De Silva N., Flint B., Frankish A., Hunt S.E., IIsley G.R., Langridge N., Loveland J.E., Martin F.J., Mudge J.M., Morales J., Perry E., Ruffier M., Tate J., Thybert D., Trevanion S.J., Cunningham F., Yates A.D., Zerbino D.R., Flicek P., 2021. Ensembl 2021. Nucleic Acids Res. 49, D884–D891. https://doi:10.1093/nar/gkaa942.

30. Satija R., Farrell J.A., Gennert D., Schier A.F., Regev A., 2015. Spatial reconstruction of single-cell gene expression data. Nat Biotechnol. 33, 495–502. https://doi:10.1038/nbt.3192

31. McCarthy D.J., Campbell K.R., Lun A.T., Wills Q.F., 2017. Scater: pre-processing, quality control, normalization and visualization of single-cell RNA-seq data in R. Bioinformatics 33, 1179–1186. https://doi:10.1093/bioinformatics/btw777.

32. Hafemeister C., Satija R., 2019. Normalization and variance stabilization of single-cell RNA-seq data using regularized negative binomial regression. Genome Biol. 20, 296. https://doi:10.1186/s13059-019-1874-1.

33. Ahlmann-Eltze C., Huber W., 2021. glmGamPoi: fitting Gamma-Poisson generalized linear models on single cell count data. Bioinformatics 36, 5701–5702. doi: 10.1093/bioinformatics/btaa1009.

34. Aran D., Looney A.P., Liu L., Wu E., Fong V., Hsu A., Chak S., Naikawadi R.P., Wolters P.J., Abate A.R., Butte A.J., Bhattacharya M., 2019. Reference-based analysis of lung single-cell sequencing reveals a transitional profibrotic macrophage. Nat Immunol. 20, 163–172. https://doi:10.1038/s41590-018-0276-y.

35. Benayoun B.A., Pollina E.A., Singh P.P., Mahmoudi S., Harel I., Casey K.M., Dulken B.W., Kundaje A., Brunet A., 2019. Remodeling of epigenome and transcriptome landscapes with aging in mice reveals widespread induction of inflammatory responses. Genome Res. 29, 697–709. https://doi:10.1101/gr.240093.118.

